# Reflected Inline Detection in Epi Oblique Plane Microscopy

**DOI:** 10.1101/2025.05.30.657110

**Authors:** Md Nasful Huda Prince, Wishwa Herath, Balasubramanian Chellammal Muthubharathi, Nikhil Sain, Chitra Shaji, Md Rafiqul Islam Rupam, Aadil Qadir Bhat, Adil R. Wani, Mubarak Hussain Syed, Tae-Hyung Kim, Mark C. Walker, Olga Ponomarova, Tonmoy Chakraborty

## Abstract

Epi oblique illumination selective plane microscopy (OPM) facilitates open-top, straightforward sample mounting, enabling high-resolution and high-speed imaging without perturbing biological specimens. However, current OPM systems typically require three separate microscopes equipped with specialized objectives. This configuration leads to complex and costly setups, significant long-term drift during imaging, reduced numerical aperture, and limited flexibility in adjusting the illumination tilt angle. We introduce Reflected Inline Detection in Epi-Oblique Plane Microscopy (RIDE-OPM), a novel, simplified, and cost-effective design that removes the necessity for a separate inline tertiary microscope and specialized objectives. Our tilt-angle-independent approach leverages the maximum NA of standard objectives, ensuring optimal illumination across the entire field of view without compromising imaging speed or quality, while significantly reducing long-term system drift. The compact and stable RIDE-OPM platform demonstrates robust, unsupervised long-term imaging capabilities. We validated this performance by successfully imaging various biological specimens, including *C. elegans*, Drosophila fly brain, human and mouse epithelial cells, and live *E. coli*, underscoring its suitability for advanced long-term biological imaging applications.

## Introduction

Light-sheet fluorescence microscopy (LSFM)^1^ has already demonstrated significant advantages over confocal and wide-field imaging, offering superior optical sectioning, gentle illumination, and faster imaging. However, the orthogonal arrangement of the illumination and detection pathways introduces two major challenges in the imaging system. First, this geometry often necessitates a specialized and complex sample mounting platform to exploit LSFM’s benefits. Various approaches have been explored to simplify sample mounting, including an inverted mounting platform that involves flipping the entire optical table—though this increases system complexity^2^—or an open-top geometry that positions both objectives beneath the sample stage^3^. Second, the orthogonal setup imposes limitations on the numerical aperture (NA) of the objective, which depends on its shape and working distance^4^. Some studies have attempted to address this by using customized illumination and detection objective pairs, but these solutions are not broadly applicable^2^. Furthermore, most of these methods require either moving the specimen or synchronously scanning the detection objective with light-sheet (LS) for 3D imaging, adding additional constraints in achieving high-resolution, fast imaging^5^. As a result, conventional LSFM with an orthogonal geometry is not well suited for high-resolution, rapid volumetric imaging.

Epi oblique illumination LS microscopy, on the other hand, offers the flexibility to utilize high-NA objectives, while eliminating the need for specialized sample mounting procedures^6^. Inspired by the oblique illumination technique used in highly inclined and laminated optical sheet (HILO) microscopy^7^, C. Dunsby and colleagues introduced the epi oblique illumination selective plane microscopy (OPM) technique^8^. Due to its unique advantages, OPM has gained attention within the imaging community and was implemented for scanning imaging^9^, high-speed volumetric imaging^10–13^, single-molecule super-resolution imaging^14–16^, structured illumination microscopy (SIM) for super-resolution imaging^17^, hyperspectral imaging of Raman microspectrometry^18^, and mesoscopic imaging for large specimen^19–23^. Continued refinements of OPM led to its current state-of-the-art implementation, which arguably is the fastest growing LS methodology^24^. Currently, the prevalent OPM designs involve three microscopes (**Fig. 1a**). The primary microscope generates a thin, inclined LS to illuminate the specimen as well as collects the emitted fluorescence. Primary and secondary microscopes together create a remote focusing arrangement that reconstructs a virtual 3D volume of the specimen. It is worth mentioning that remote focusing^25^ itself has been widely adopted in various imaging systems, including spinning disc^26^, multiphoton^27^, axially swept LS^28,29^, and hybrid open-top microscopy^3^ setups. Through appropriate pupil matching of the primary (O_I_) and secondary (O_II_) objectives, OPM forms a tilted virtual image plane of the sample after O_II_, ensuring identical lateral and axial magnification^30,31^. The images of any specific 1D lines in the sample’s xz’ plane are generated after the secondary objective. A tertiary microscope, positioned orthogonally to this virtual image plane, captures the xz’ plane at the camera frame with all possible 1D lines as shown in the tilted sample plane (**Fig. 1a**). By scanning the specimen along the y-axis, a full 3D volume can be reconstructed. A deskewing algorithm then rotates this volume to generate an accurate 3D representation of the specimen^32^. Since OPM removes the constraint of orthogonal illumination-detection geometry and enables 3D imaging with laser scanning, OPM offers a promising alternative for achieving high-resolution and fast imaging in LS based microscopy. However, as discussed later, its tilted tertiary microscope configuration complicates alignment and increases system cost due to the need for a customized tertiary objective, which also leads to a loss of NA at the secondary-tertiary microscope interface.

**Fig. 1.**
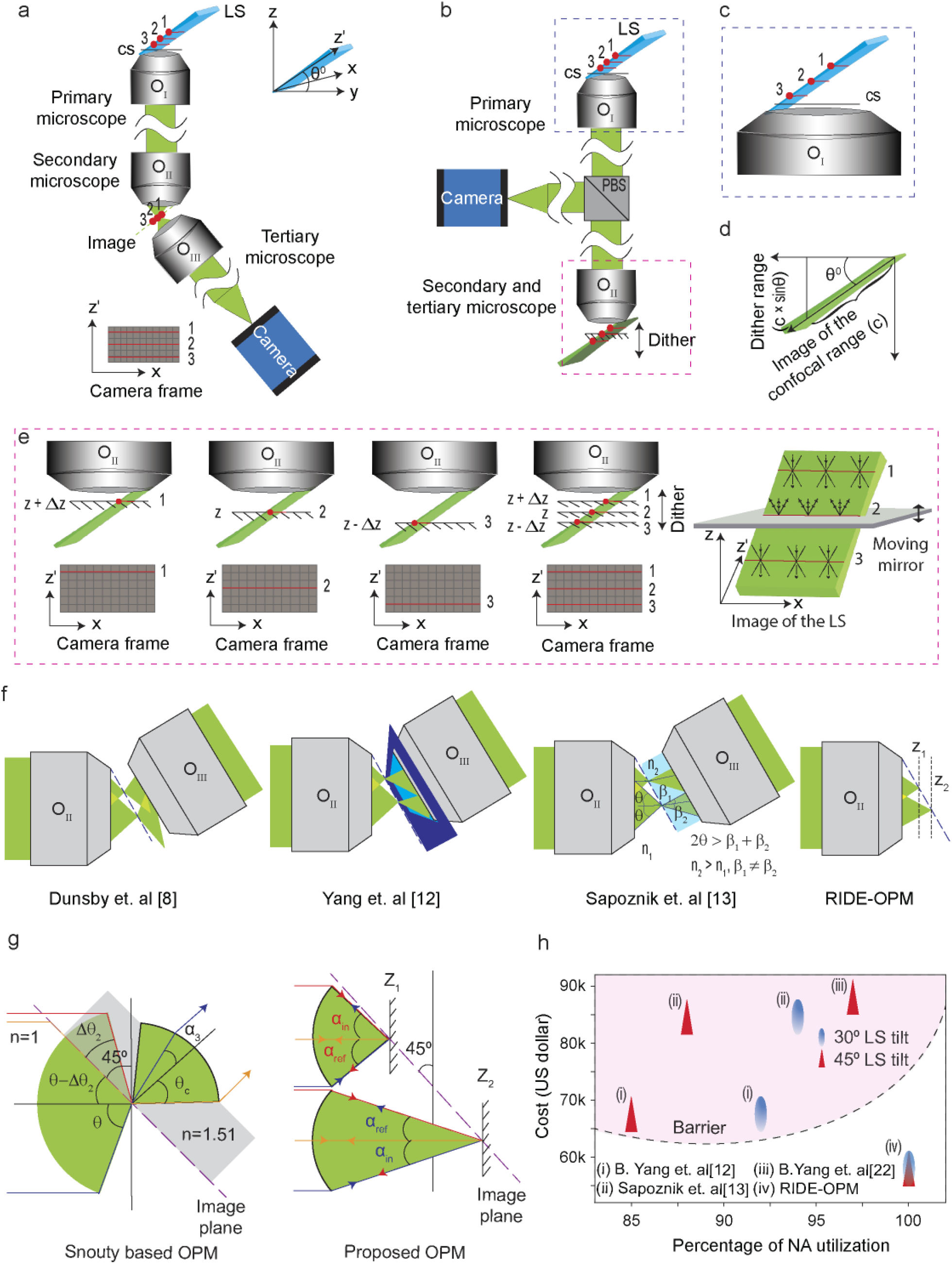
Optical principle of epi oblique plane microscope. **a**, Conventional OPM setup consisting of three microscopes: primary, secondary, and tertiary. A tilted LS illuminates the specimen, and the image of the xz’ sample plane is reconstructed after O_II_ using a pupil-matched remote focusing system. The tertiary microscope, positioned orthogonally to this virtual image plane, captures multiple 1D lines within the xz’ plane to form a 2D image. **b**, The proposed RIDE-OPM setup eliminates the tertiary microscope, with the secondary microscope serving both roles. A dithering mechanism is employed after the secondary microscope to reflect the entire image plane back to the camera. **c**, A zoomed-in view of O_I_ and the tilted image of the LS of **b**, illustrating multiple 1D lines distributed along the z-axis within the 2D plane. **d**, The minimum dither range is determined by multiplying the image length of the LS confocal range by the sine of the LS tilt angle. **e**, A static mirror at different z-positions captures individual 1D lines (shown in the rightmost), while a mirror dithering at twice the camera speed reconstructs the full xz’ plane. **f**, Fluorescence collection at the O_II_ - O_III_ interface. High-refractive-index (RI) media (e.g., water) in customized setups, and later the snouty objective, were used in conventional OPM to capture the entire light cone, preventing signal loss. The proposed RIDE-OPM achieves full fluorescence collection by maintaining identical incoming and collection light paths. **g**, Snouty-based OPM loses NA with increasing LS tilt (**Supplementary Fig. 1**), while RIDE-OPM is tilt-independent, ensuring full NA utilization (**Supplementary Fig. 2**). **h**, The proposed RIDE-OPM system eliminates cost and NA utilization barriers associated with LS tilt, making high-NA OPM implementations more accessible and efficient. LS, light-sheet; cs, coverslip; O, objective; PBS, polarizing beam splitter.

Recognizing the limitations associated with the tertiary objective, several prior studies have explored inline OPM configurations for specimen imaging (e.g., obSTORM^14^ and dOPM^33,34^). In single-molecule oblique-plane microscopy (obSTORM), a tilted 2D plane was imaged; however, this approach suffers from significant fluorescence loss, leading to a reduction in effective NA and consequently limiting the system from achieving diffraction-limited resolution. Although the incorporation of stochastic optical reconstruction microscopy (STORM) enabled super-resolution imaging, it remained restricted to two-dimensional reconstruction. In another approach, dual-view oblique plane microscopy (dOPM), a complex prism–mirror configuration was employed to reflect the fluorescence signal and de-scan the light-sheet plane onto the camera. While dOPM eliminates the need for a separate tertiary objective, it introduces substantial complexity in alignment and requires precise mechanical translation of the prism assembly. This design not only reduces the effective fluorescence collection efficiency but also significantly compromises imaging speed and introduces additional computational steps for image fusion. In a separate approach, an electro-tunable lens (ETL) has been employed to refocus different one-dimensional sections of the light-sheet^35^. Although this method captures the full fluorescence collected by O_I_, aberrations introduced by the ETL degrade image quality and result in reduced spatial resolution. Additionally, the need for tight synchronization between the ETL modulation and the rolling shutter of the camera further limits imaging speed (**Supplementary Note 1** and **Supplementary Table 1**).

To overcome these limitations, we propose a new OPM with a simple and compact inline design for the detection path for fluorescence. We call it Reflected Inline Detection in Epi-Oblique Plane Microscopy (RIDE-OPM). Our RIDE-OPM design eliminates the necessity of a separate, and often tilted, tertiary microscope, significantly reducing costs by removing the need for an expensive, highly specialized tertiary objective while maintaining imaging performance (**Fig. 1b-1e**). The proposed setup ensures optimal light collection across the entire field of view (FOV) and preserves the NA of the secondary objective, addressing a major limitation of the current OPM design^13^. Our proposed RIDE-OPM integrates a dithering device and is optimized to achieve camera-limited imaging speed for desired FOV, ensuring an efficient and effective imaging approach.

## Results

### Instrument design considerations

A critical and recurring factor in the evolution of OPM designs has been the issue of maintaining consistent light collection and NA across the FOV. This poses a significant challenge in OPM designs, particularly at the interface between the secondary and tertiary objectives, due to its geometric constraints^36^. Several approaches have been proposed to address this issue (**Fig. 1f**). For example, Yang et al. attempted to mitigate the problem by introducing a higher refractive index (RI) medium—water with RI 1.33—between the secondary and tertiary interfaces^12^. This method improves performance by effectively compressing the detection light cone, enabling the tertiary objective (O_III_) to capture a larger angular range of light coming from O_II_. Building on this concept, Millett-Sikking et al. designed a custom glass-tipped air objective (NA∼ 1.0), commonly referred to as “Snouty”, to optimize the light collection^37^. Sapoznik et al. further validated this approach through theoretical optical ray diagrams, demonstrating that while “Snouty” somewhat alleviated the issue, the half-aperture angles of O_III_ relative to O_II_ still remained asymmetric^13^. Nonetheless, several recent OPM systems have incorporated this customized objective, showcasing its effectiveness despite its complexity and high cost.

While the specialized design of “Snouty” effectively addresses the issue of light collection at the secondary and tertiary microscope interface, it introduces another limitation—tilt-dependent performance. In OPM, the tilt of the LS plays a crucial role in determining image resolution, confocal range, and the accessible z-depth above the coverslip^13^. Ideally, a microscope should be independent of LS tilt, providing greater flexibility for users to adjust imaging parameters based on their specific needs. However, the glass piece in front of the snouty objective reduces the light collection aperture angle, leading to a loss of NA and consequently a decrease in effective resolution compared to what the objective could otherwise achieve (**Fig. 1f**). Although Sapoznik et al. demonstrated that the imaging capability of the “snouty” remains largely intact with only a minor NA loss for a 30° LS tilt, this loss becomes increasingly significant as the LS tilt angle increases (**Supplementary Fig. 1**).

In contrast, our proposed RIDE-OPM overcomes this issue by reflecting the light orthogonally to the optical axis, ensuring a symmetric aperture angle in O_III_ and capturing the entire light cone (**Supplementary Fig. 2**). As a result, RIDE-OPM guarantees complete utilization of the NA offered by the primary and the secondary objective, regardless of the LS tilt (**Fig. 1g**). Furthermore, the complexity of the OPM design, its high cost, and the tilt-dependent NA loss pose significant barriers to the widespread adoption of existing OPM platforms. RIDE-OPM design eliminates these challenges through a simpler, more cost-effective approach that removes the need for a separate tertiary objective altogether, while fully utilizing the NA of the secondary objective (**Fig. 1h**). This streamlined design enhances the imaging performance while making high-resolution, flexible imaging more accessible.

### Long-term Stability and Performance Challenges in OPM Imaging Systems

As OPM gains traction within the biomedical imaging community, significant technical hurdles associated with the tertiary objective are becoming increasingly evident. These include (a) the complexity involved in setting up the tertiary microscope at an inclined angle, (b) aberrations introduced due to this inclined geometry, and (c) the compromised long-term stability resulting from mechanical drift. For example, as pointed by Sapoznik et al. OPMs equipped with inclined tertiary objectives suffer from a time-dependent drift between the secondary and tertiary objectives^13^, which limits the system’s ability to maintain optimal performance during extended imaging sessions. This drift is primarily caused by the well-known sensitivity of microscope objectives to temperature fluctuations, where even a minor temperature fluctuation can cause axial shifts in the focal plane—a problem that becomes particularly pronounced with high NA objectives due to their inherently shallow depth of focus. Millett-Sikking et al. quantified this thermal sensitivity using a Nikon 40×/0.95 NA objective and reported an axial focal drift of approximately 1.58 µm per °C^38^. In OPM systems, both lateral and axial drifts in the secondary and tertiary objectives have been observed^39^. These drifts cause the image of the LS to shift from its original position, misaligning it from the nominal focal plane (NFP) of the tertiary objective—a critical alignment requirement for achieving sharp focus in OPM. As a result, the point spread function (PSF) deteriorates over time, becoming broadened or defocused (**Supplementary Fig. 3**). While one common mitigation strategy is to mount the tertiary objective on a piezo actuator to actively correct for drift, this adds complexity and imposes limits on the microscope’s ability to reliably capture time-sensitive biological processes^40,41^. These documented issues demonstrate that precise alignment and consistent performance of a separate, inclined, tertiary microscope in OPM systems are inherently difficult to maintain, requiring frequent manual intervention, meticulous alignment procedures, and often costly active correction strategies.

### Microscope design

The main principle underlying RIDE-OPM involves understanding that the remote focusing forms an image of the tilted xz’ LS plane, in accordance with the LS angle. This secondary image can be visualized to be composed of numerous individual points, each corresponding to a distinct cone of light (**Fig. 1e**). The important thing to realize here is that, although the image of the xz’ LS is tilted, the fluorescent light that makes up this image still propagates straight along the optic axis of O_II_. When a flat mirror is placed orthogonally to the optic axis, the tilted image plane intersects with the mirror plane along a one-dimensional (1D) line (shown as red lines in **Fig. 1e**). It is because of this orthogonal reflection, light at this intersection reflects directly back along its original path, efficiently capturing the reflected fluorescence (**Supplementary Fig. 4** and **Supplementary Note 2**). We therefore make use of an optical isolator in the detection arm (**Fig. 1b**), prior to O_II_, such that the reflected fluorescence light can be extracted and put on an sCMOS camera for imaging. This however means, with a static mirror, the sCMOS camera captures only an individual 1D line of the image plane.

To fully capture the tilted image formed by O_II_, RIDE-OPM uses a mechanical dithering mechanism that moves a mirror along the optical axis. By adjusting the axial position of the mirror, different 1D lines across the virtual tilted plane can be selectively reflected and recorded. This principle enables stepwise acquisition of the entire 2D image plane through sequential 1D line reflections. However, to achieve real-time capture of the full tilted image within a single exposure time, we implemented rapid axial dithering of the mirror, oscillating it back and forth at a frequency at least twice the rate of the camera exposure time. This ensures that the mirror traverses the entire depth of the light sheet’s (LS’s) confocal range within each exposure cycle. The net effect is a time-averaged integration of all 1D slices, forming a complete and continuous image of the tilted plane (**Supplementary Note 3**). Additionally, the dither step size adheres to the Nyquist criterion, essential for accurate reconstruction of continuous image features (**Supplementary Fig. 5**).

From the perspective of long-term imaging and system simplicity, RIDE-OPM eliminates the need for a tertiary objective, resulting in a more compact system footprint and significantly reducing the dominant sources of drift found in conventional OPM configurations. While the O_II_ may still experience time-dependent drift, the reflective mirror in the RIDE-OPM system can dynamically compensate for this movement across its operational range, maintaining alignment and focus (**Supplementary Note 4**). This makes RIDE-OPM a robust and reliable platform for long-term unsupervised imaging. Additionally, the system’s inline optical configuration eliminates tilt-induced aberrations, further enhancing image stability and fidelity over extended acquisition periods.

### Microscope characterization

We operated the RIDE-OPM system in two distinct configurations, scale 1 and scale 2, utilizing different combinations of O_I_, O_II_, and relay lenses. To evaluate the system’s tilt-independent performance within the range of 30°-45° tilt, scale 1 was tested at both 30° and 45° LS tilt angles. For each condition, we adjusted the effective illumination NA using an adjustable slit to achieve the desired confocal range and thereby match the imaging depth requirements of the specimen. Specifically, scale 1 was operated at illumination NAs of 0.30 and 0.36. Scale 2 was configured to provide a longer confocal range and greater imaging depth, with a slight trade-off in axial resolution (NA 0.16). A conjugated high-speed galvo mirror was used to scan the LS along the y-axis, generating a lateral image (**Fig. 2a**). During post-processing, the deskewing algorithm preserved the LS extent along the x-axis while rotating and resampling the y and z dimensions, resulting in final image volumes that varied based on LS tilt and system scale (**Table 1, Fig. 2b** and **Supplementary Fig. 6**). Compared to the theoretical axial depth limits associated with the 80% Strehl ratio for remote focusing, 53 µm for scale 1 and 162 µm for scale 2, our measured imaging depths were relatively shallow (4–9 µm for scale 1, 30 µm for scale 2), thereby ensuring uniform optical performance throughout the entire z-range (**Supplementary Fig. 7**).

**Fig. 2.**
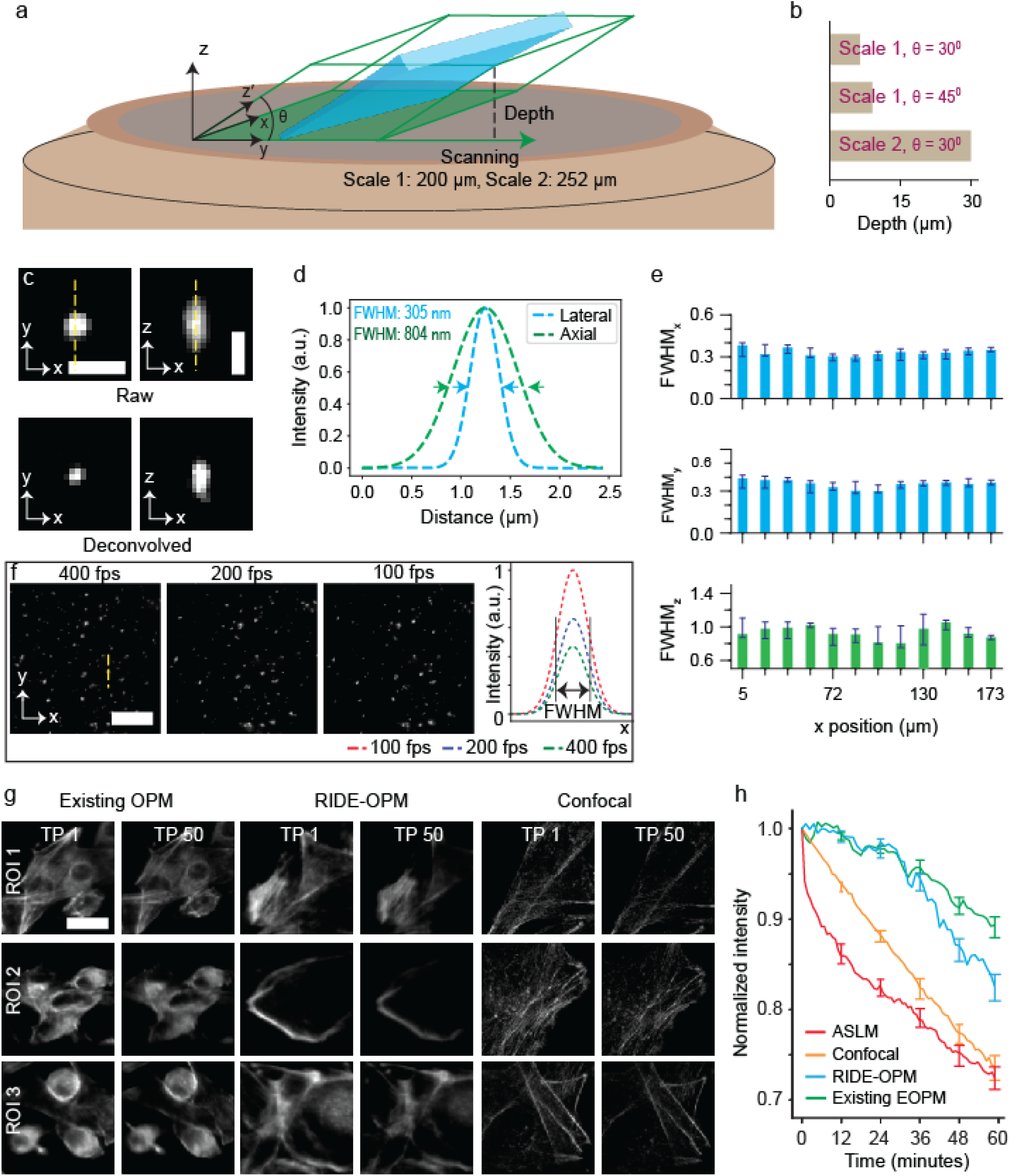
Microscope characterization. **a**, Imaging volume. Considering z is the direction of the optic axis, a LS was created at an angle θ with respect to the xy plane to illuminate the specimen, where x is the LS length and z’ is the direction of propagation. Camera captured the xz’ plane with a FOV of x × y × z’ where z’ is the confocal range of the LS (**Table 1**). 3D volumetric image was acquired by laser scanning along the y axis, resulting in a volume with expected depth. **b**, Imaging depths achieved with scale 1 (0.30 NA) at different LS tilt angles and with scale 2 demonstrate the inherent trade-off between tilt angle, resolution, and imaging depth. **c**, Orthogonal representation of both raw and Richardson-Lucy deconvolved PSF, imaged 100 nm of fluorescent beads embedded in glycerol. **d**, Resolution was determined from the Gaussian-fitted line profile. **e**, Resolution measurements across the entire FOV demonstrated consistent imaging performance throughout. Panels **c, d** and **e** correspond to scale 1 with 30° LS tilt. **f**, A zoomed-in view of a randomly selected small region from the entire FOV for scale 2 (**Supplementary Fig. 9**), showing 500 nm fluorescent beads embedded in glycerol imaged at various speeds. Line profiles of a random bead demonstrate consistent image quality with varying intensities. **g-h**, Comparative photobleaching rates of existing OPM, RIDE-OPM, confocal, and ASLM microscopes in MDA-MB-231 human triple-negative breast cancer cells, stained for F-actin with phalloidin–Alexa Fluor 647. Fluorophore bleaching was measured across multiple ROIs over 50 time points acquired within a one-hour period (**g**). The images acquired from both RIDE-OPM and OPM are presented in their raw, unprocessed form. Photobleaching rates for existing OPM and RIDE-OPM were comparable, with both showing significantly gentler performance compared to confocal and ASLM systems, where the bleaching rate was approximately 2× faster (**h**). Scale bars, 1 µm (**c**), 25 µm (**f**), 10 µm (**g**). fps, frames per second; FWHM, full width at half maximum.

**Table 1.** Microscope characterization for various scales and tilt angles.

| Type | LS tilt angle | Illumination NA | FOV (xz') ($\mu\text{m} \times \mu\text{m}$) | Depth ($\mu\text{m}$) | FOV (xy) ($\mu\text{m} \times \mu\text{m}$) | Resolution | | |
| --- | --- | --- | --- | --- | --- | --- | --- | --- |
|  |  |  |  |  |  | Lateral (n = 30) |  | Axial (n = 30) |
|  |  |  |  |  |  | x-axes (nm) | y-axes (nm) |  |
| Scale 1 | 30° | 0.30 | 173 × 12 | 6 | 173 × 200 | 318 ± 6<br>216 ± 8* | 302 ± 8<br>206 ± 7* | 1 ± 0.04 $\mu\text{m}$<br>848 ± 6 nm* |
| | | 0.36 | 173 × 8.5 | 4.2 | | | | 802 ± 7 nm<br>605 ± 7 nm* |
| | 45° | 0.30 | 173 × 12 | 9 | | 322 ± 8<br>206 ± 5* | 306 ± 7<br>204 ± 6* | 1 ± 0.04 $\mu\text{m}$<br>848 ± 6 nm* |
| | | 0.36 | 173 × 8.5 | 6 | | | | 802 ± 7 nm<br>605 ± 7 nm* |
| Scale 2 | 30° | 0.16 | 200 × 60 | 30 | 200 × 252 | 446 ± 10<br>341 ± 18* | 455 ± 8<br>352 ± 14* | 1.81 ± 0.06 $\mu\text{m}$<br>1.63 ± 0.05 $\mu\text{m}$* |

\* RL deconvolution

To experimentally characterize the resolution of the system, we imaged 100 nm fluorescent beads embedded in glycerol under 488 nm laser excitation. The full width at half maximum (FWHM) of the bead intensity profiles was used as a resolution metric (**Fig. 2c–2d**). As expected, resolution remained consistent across the entire FOV (**Fig. 2e, Supplementary Fig. 8**). We observed slight PSF asymmetry, which can be attributed to several factors, including inherent optical astigmatism, sample-dependent wavefront distortions, refractive index (RI) mismatch, and residual aberrations in the remote focusing system. Applying Richardson–Lucy deconvolution with a predefined experimental PSF further enhanced spatial resolution (**Table 1**). Finally, with the current camera and mechanical mirror dithering setup, we achieved camera-limited imaging speeds even for the 30 µm axial depth of scale 2 (**Fig. 2f, Supplementary Fig. 9**). Collectively, these results demonstrate that RIDE-OPM enables high-resolution, fast 3D volumetric imaging with stable and consistent performance across a range of LS tilt angles and field sizes without any trade-off.

### Photobleaching comparison

We experimentally evaluated the photobleaching rates of various imaging systems over a continuous one-hour acquisition period. The performance of the RIDE-OPM system was compared against an in-house built state-of-the-art OPM system, a commercial confocal microscope, and an axially swept light-sheet microscopy (ASLM) system (see methods for system details) (**Fig. 2g–2h, Supplementary Fig. 10-11**). For this comparison, we imaged human breast adenocarcinoma (MDA-MB-231) cells stained with Alexa Fluor 647 targeting F-actin. To ensure a fair comparison, we fixed the starting image intensity across all systems by adjusting the laser power for each setup. The ASLM system began with slightly lower baseline intensity due to system limitations. All datasets were acquired with a camera exposure time of 100 ms, and volumetric imaging in the RIDE-OPM system was performed with a 280 nm step size. We anticipated that several factors—including the presence of a thick glass element in the snouty objective, fluorescence loss at the O_II_-O_III_ interface in conventional OPM, and the inherently gentle illumination characteristics of OPM systems—could differentially affect photobleaching behavior between RIDE-OPM and conventional OPM. Consequently, a difference in intensity decay between the two systems was observed over the one-hour imaging period. Both systems exhibited nearly identical photobleaching rates during the initial phase. After this period, RIDE-OPM showed a marginally faster decline in fluorescence intensity. Nevertheless, both OPM configurations significantly outperformed the confocal and ASLM platforms, which exhibited photobleaching rates at least twice as high as that of RIDE-OPM.

This result highlights the gentler illumination of RIDE-OPM compared to widely used modalities such as confocal and ASLM. The experiment was repeated for five times for each system, and all trials consistently exhibited the same bleaching trends (**Fig. 2h**).

### Imaging of multi-ROI and tracking *C. elegans* larval development

*C. elegans* research often involves quantification of phenotypes with varying degrees of penetrance (the proportion of individuals with a specific genetic variant who actually display the associated phenotype). Observing multiple individual animals in parallel on a single slide not only speeds up the analysis but also makes it feasible to quantitatively study inter-individual differences within a population. We evaluated RIDE-OPM’s capability to automatically image multiple regions of interest (ROIs) (**Fig. 3a**). We used low irradiance to navigate through the sample and selected 15 distinct ROIs with varying imaging relevance. These were then sequentially imaged through automated stage movements under optimal irradiance conditions (**Fig. 3b–3d**), enabling identification of healthy ROIs exhibiting features of interest. To assess long-term imaging performance, we tracked the developmental differences between the wild type (**Fig. 3e**) and the *idh-1neo* mutant (**Fig. 3f**) *C. elegans*, which carries a neomorphic *idh-1* gene mutation^42^ for over ∼13 hours. The healthy wild type embryo moves continuously through the final stage of embryogenesis and begins to hatch from its eggshell (**Supplementary Movie 1**). In contrast, the *idh-1neo* mutant embryo exhibits reduced movement, eventually ceases all activity, and fails to hatch (**Supplementary Movie 2**). This underscores the system’s robustness for long-term, unsupervised imaging, with its inline design further enhancing stability and image fidelity. Furthermore, the system consistently delivers adequate signal-to-noise ratios with minimal photodamage, affirming its suitability for sensitive biological imaging tasks.

**Fig. 3.**
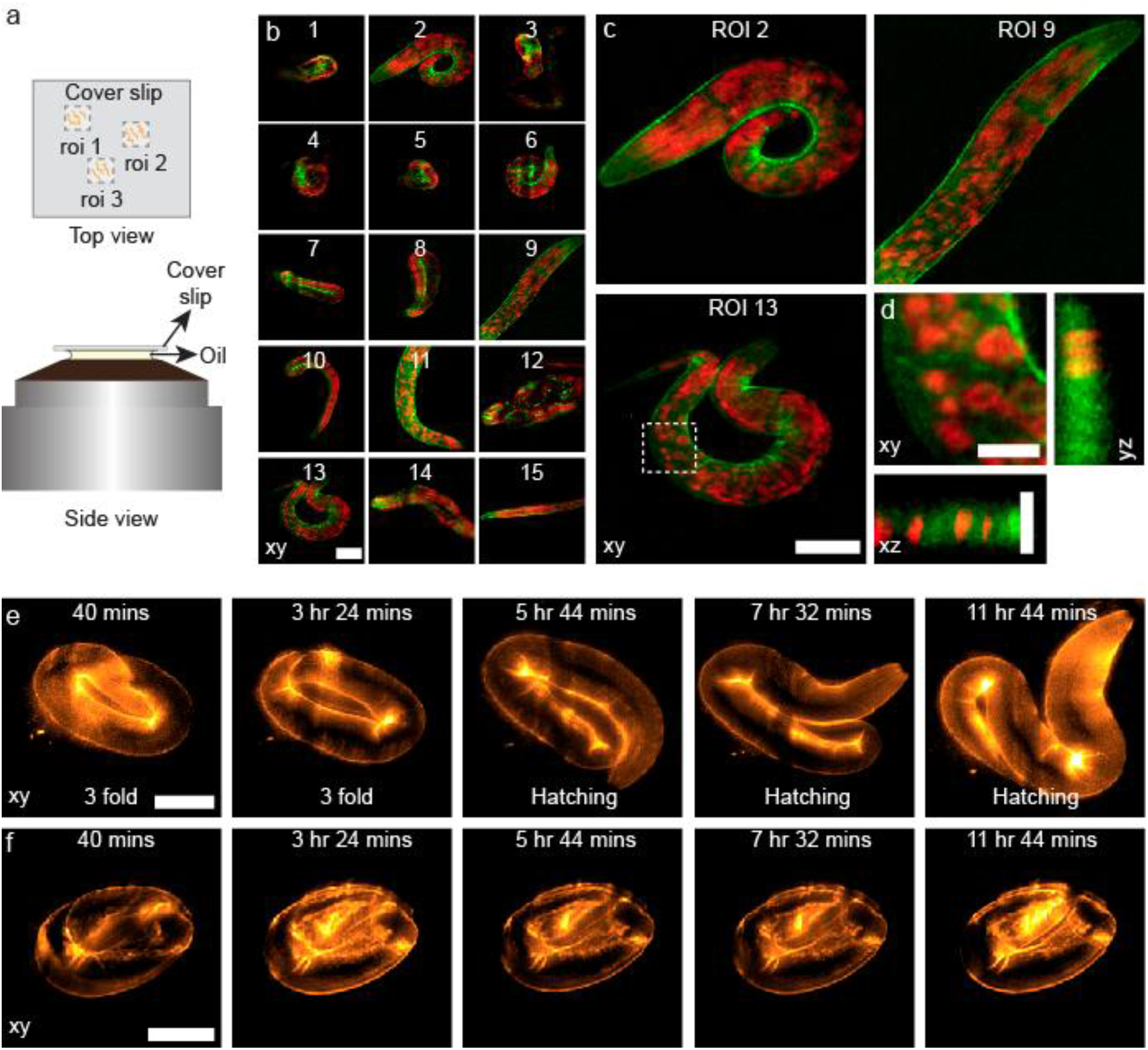
Unsupervised, multi-ROI imaging and long-term developmental tracking of *C. elegans* embryo and L1 larvae. **a**, Top and side views of the imaging setup with the mounted specimen. Based on the specimen’s growth pattern, concentration, and orientation, multiple ROIs can be selected for imaging and analysis. **b**, Automated sequential imaging of 15 ROIs, each representing a *C. elegans* L1 larva with *idh-1* neomorphic mutation (*idh-1neo*^42^) expressing endogenously tagged cuticle collagen SQT-3::mNG (green), NucSpot staining highlights nuclei (red). **c**, Zoomed-in view of three healthy ROIs. **d**, Orthogonal views of the selected region, marked by the white dashed box in ROI 13 of **c**, confirm effective optical sectioning along the z-axis. The xz and yz projections represent maximum intensity projections (MIPs) of only three consecutive slices. **e-f**, Time-lapse imaging of developmental progression in wild-type (**e**) and *idh-1neo* (**f**) *C. elegans* embryos expressing the tagged cuticle collagen SQT-3::mNG. Imaging was initiated at the 3-fold embryonic stage. Images were acquired every 4 minutes over a 13-hour period (**Supplementary Movie 1-2**). Scale bars, 20 µm (**b-c, e-f**), 5 µm (**d**). roi, regions of interest.

### Mesoscopic imaging

Combining whole-animal imaging with fluorescent reporters such as *col-19::GFP*, a stage- and tissue-specific marker, enables the assessment of alterations in the timing of developmental events (heterochronic phenotypes)^43,44^. We assessed the microscope’s ability to extend its FOV by imaging whole millimeter-scale *C. elegans* (**Fig. 4a–4b**) and Drosophila fly brain using a tiling approach via scale 2 to cover a longer depth above the cover slip. Given the limited FOV of 200 µm × 250 µm in the xy-plane, we acquired ten tiles (five in x, two in y) with 10% overlap in all directions to capture the full *C. elegans* organism. The zoomed-in views reveal fine structural details in both lateral and axial planes (**Fig. 4c**). Each tile spanned 200 µm × 250 µm × 30 µm (**Fig. 4d–4e**), with the 30 µm z-depth sufficient to cover the full body thickness of adult *C. elegans*. We applied a similar tiling strategy to image an entire adult Drosophila brain, positioned between a coverslip and cover glass. The brain was captured in six overlapping tiles. The image of R84C10 dorsal fan-shaped body neurons, labeled with GFP, clearly shows cell bodies, dendrites, and axons in three dimensions (**Fig. 4f–4g**). A line profile along the brown dotted line in **Fig. 4f** highlights the ability to distinguish two lobes of the dendritic projections, marked by the red arrowhead in the lateral inset (**Fig. 4h**). A similar profile along the red dotted z-line in **Fig. 4g** shows a ∼2 µm FWHM based on Gaussian fitting (**Fig. 4i**).

**Fig. 4.**
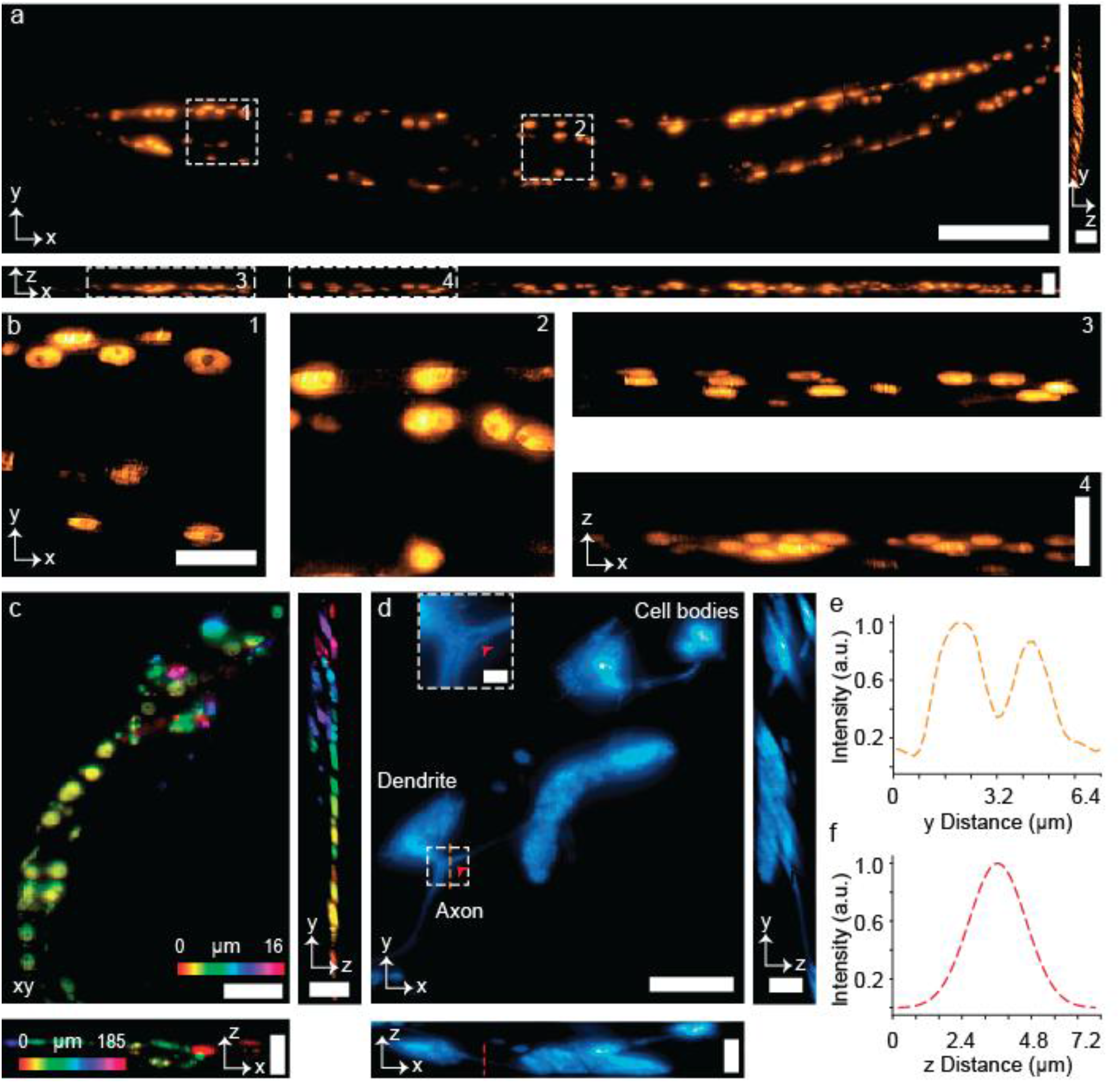
Mesoscopic imaging for extended field-of-view and structural resolution. **a-b**, Tiled reconstruction of a ∼ millimeter-sized *C. elegans*, expressing adult-specific *col-19::GFP* transgene which marks hypodermal nuclei. Lateral and axial views show the full body, with zoomed-in regions (dotted boxes) highlighting the expression of a transgenic protein (**b**). **c**, A single image tile spanning a 30 µm depth shows both lateral and axial views, demonstrating the resolution within one tile. **d**, Orthogonal views of R84C10 dorsal fan-shaped body neurons labelled with GFP in an adult Drosophila brain. Insets show zoomed-in dendritic projections. The imaged volume was 200 µm × 252 µm × 30 µm. **e-f**, Line profile over the brown and red line corresponding to the lateral and axial view of **d**, respectively. The lateral line profile (**e**) highlights distinct dendritic projections (red arrowhead), while the axial profile (**f**) shows a Gaussian-fitted FWHM of ∼2 µm. Scale bars, 100 µm (xy) and 20 µm (xz, yz) (**a**), 20 µm (**b**), 30 µm (xy) and 20 µm (xz, yz) (**c**), 50 µm (xy) and 20 µm (xz, yz) (**d**), 5 µm (inset of **d**).

### Tilt-independent multi-channel imaging in the RIDE-OPM System

We further investigated the tilt-independent performance of the RIDE-OPM system within the 30°–45° range, the most commonly used oblique illumination angles in OPM system. This range inherently involves a trade-off between LS tilt angle, spatial resolution, and imaging depth above the coverslip (**Supplementary Note 4**). Ideally, a system should allow seamless switching between different tilt angles without compromising imaging performance. However, this remains technically challenging for conventional OPM systems. In conventional OPMs, changing the LS tilt angle alters the orientation of the virtual image (xz′ plane) formed after O_II_. To maintain optimal imaging, the narrow nominal focal plane (NFP) of O_III_ must remain aligned with this tilted image plane. Adjusting for such alignment requires precise repositioning of the entire tertiary optical assembly, including O_III_, the tube lens, and the camera, both rotationally and translationally (**Fig. 5a**). This complexity typically limits conventional OPM systems to operate at a single, fixed tilt angle.

**Fig. 5.**
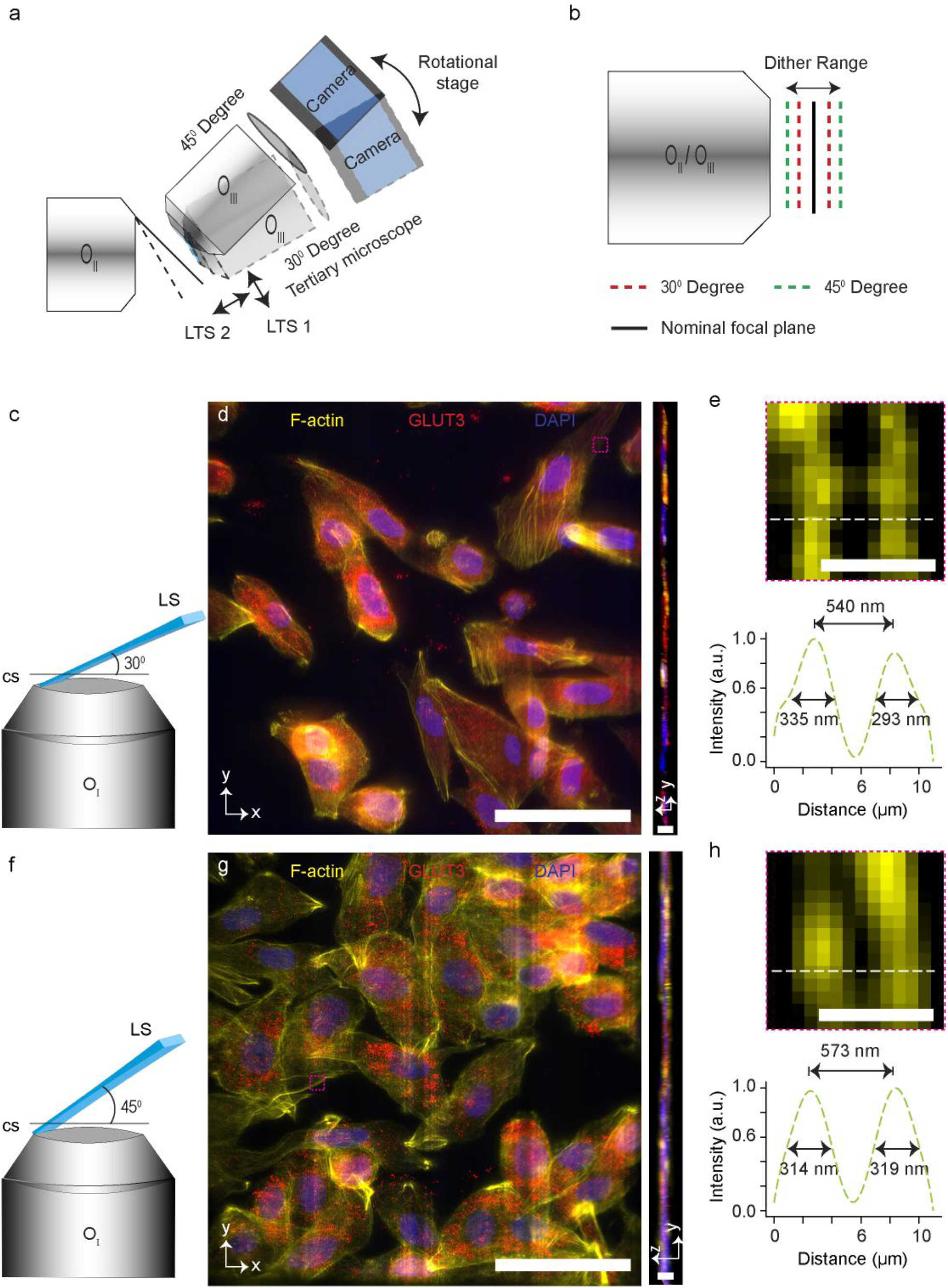
Tilt-independent multi-color imaging. **a-b**, Comparison of physical alignment requirements for switching the LS tilt angle between conventional OPM (**a**) and RIDE-OPM (**b**). In conventional OPM systems, changing the LS tilt angle requires realigning the virtual image formed after O_II_ with the NFP of the tertiary objective. This process demands repositioning the entire tertiary microscope, including O_III_, tube lens, and camera, not only through rotation but also with precise two-dimensional translation. In contrast, RIDE-OPM eliminates this complexity by not requiring any physical realignment to accommodate various LS tilt angle. Switching the LS tilt angle only requires adjusting the axial dithering range, without the need to reposition any optical elements. LST, linear translation stage. **c-h**, MDA-MB-231 (human triple-negative breast cancer) cell, imaged with 30° (**c-e**) and 45° (**f-h**) LS tilt angles. The cell was stained for F-actin (yellow), GLUT3 (red), β-tubulin (green), and DAPI (blue; nuclei). F-actin was visualized using phalloidin-Alexa Fluor 647. GLUT3 staining was performed using a primary antibody (anti-GLUT3, 1:200) and Alexa Fluor 561-conjugated secondary antibody. β-tubulin was labelled with an anti-β-tubulin antibody (1:500) and Alexa Fluor 488 secondary antibody. Expanded views of the magenta boxed regions from **d** (**e**) and from **g** (**h**) of F-actin (top), with corresponding intensity profiles across the white dashed lines. The profiles clearly resolve two actin fibers separated by 540 nm and 573 nm, respectively. Scale bars, 50 µm (xy of **d**, and **g**), 6 µm (yz of **d**, and **g**), 1 µm (**e, h**). LTS, linear translation stage.

In contrast, RIDE-OPM eliminates the need for a tertiary objective, thereby removing the constraints of this mechanical realignment. The system can switch between different LS tilt angles by simply adjusting the dithering range of the mirror through software, without any physical reconfiguration (**Fig. 5b**). This allows users to select optimal imaging conditions based on sample geometry and depth requirements, without compromising resolution or signal quality.

To demonstrate the tilt-independent, multicolor imaging capability of RIDE-OPM, we imaged fixed samples from three different cell lines using two optical scales. Scale 1 was used to image human breast adenocarcinoma cells (MDA-MB-231) (**Fig. 5** and **Supplementary Fig. 12-13**) and murine mammary adenocarcinoma cells (EO771) (**Supplementary Fig. 14**), while scale 2 was used for imaging non-tumorigenic human mammary epithelial cells (MCF10A) (**Supplementary Fig. 15**). We targeted key cellular structures including F-actin, β-tubulin, GLUT3 (glucose transporter 3), and nuclei. F-actin was labeled with Alexa Fluor 647, β-tubulin with Alexa Fluor 488, GLUT3 with Alexa Fluor 561, and nuclei with DAPI. RIDE-OPM achieved robust multichannel imaging with strong signal detection across all four channels at both 30° (**Fig. 5c–5e**) and 45° (**Fig. 5f–5h**) oblique illumination angles.

The system revealed distinct spatial distributions of cytoskeletal and membrane-associated markers across all cell types. Notably, two closely spaced actin fibers, separated by approximately 550 nm center-to-center, were clearly resolved in both tilt conditions, with an apparent width of ∼300 nm for each microtubule, demonstrating the system’s high spatial resolution under varying LS tilt configurations (**Fig. 5e, Fig. 5h**).

### High-speed imaging of live specimens

We evaluated the imaging speed of the microscope by capturing live *E. coli* cells containing superfolder green fluorescent protein (sfGFP) and *C. elegans* embryos at 200 frames per second (fps) (5 ms of camera exposure). sfGFP-expressing *E. coli* cells, freely moving in phosphate-buffered saline (PBS) on a coverslip, were imaged over a 130 µm axial scan range with a step size of 650 nm. Each 200 µm × 130 µm × 15 µm volume was acquired in one second, and 200 consecutive time points were captured to monitor bacterial motion. A maximum intensity projection (MIP) from a single time point shows the position of *E. coli* cell at that instant (**Fig. 6a**), while a MIP of all 200 time points illustrates their 2D trajectories over a 200-second period (**Supplementary Movie 3**). Representative track of *E. coli* over a 5-second interval highlights variations in movement (**Fig. 6b**). This high-speed imaging enabled quantitative tracking of *E. coli* motion, revealing variability in speed and distance traveled. Some bacteria remained stationary, while others showed distinct motion patterns. Based on maximum speed, *E. coli* was categorized into three groups, and similarly grouped again by total 2D distance travelled (**Fig. 6c–6d**). Segmentation from a single time point (**Fig. 6e**) allowed us to measure the 2D area occupied by the bacteria (**Fig. 6f**). We also imaged a *C. elegans* L1 larva under the same high-speed conditions, successfully tracking its movement over a 200-second duration (**Fig. 6g**). High-speed imaging enables the observation and capture of a range of movements in live animals, which could potentially be used to analyze diverse behavioral and neurological phenotypes (**Supplementary Movie 4**).

**Fig. 6.**
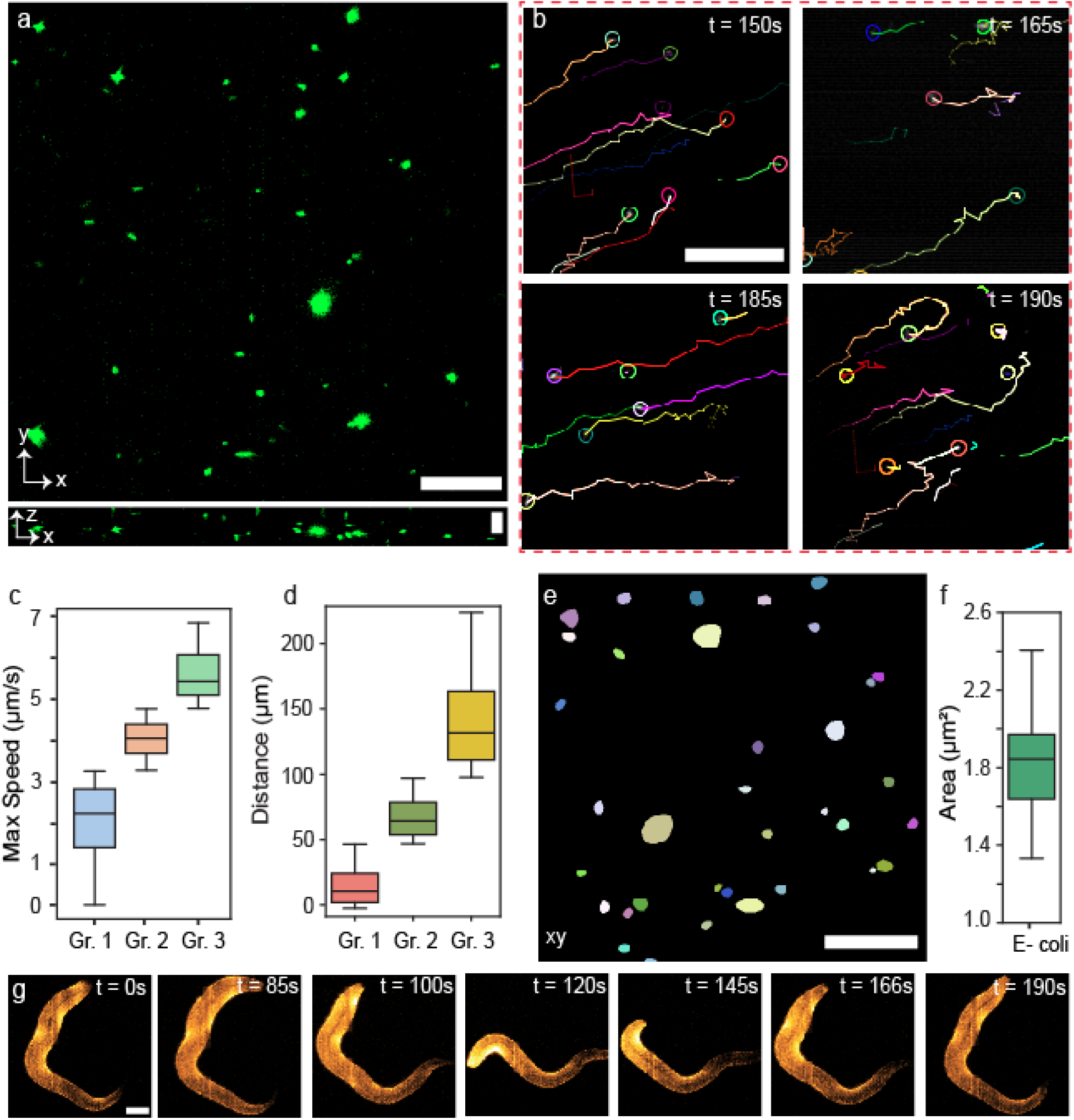
High-speed imaging and motion analysis of *E. coli* and *C. elegans* embryos. **a**, Maximum intensity projection (MIP) for in both lateral and axial views of a single time point showing sfGFP expressing *E. coli* imaged at 200 fps. **b**, High-speed imaging enables tracking of *E. coli* movement caused by the presence of flagella. 2D tracks over a 5-second period from different regions and time points illustrate directional motion. **c-d**, Motion analysis of *E. coli* based on speed (**c**) and distance travelled (**d**). Bacteria were grouped into three categories in each case, revealing heterogeneous movement behaviors. **e-f**, Segmentation of *E. coli* from a 2D MIP (***e***) enabled measurement of the 2D area occupied by each bacterium (**f**). **g**, The same 200 fps imaging setup was used to track live, moving *idh-1neo C. elegans* in L1 larval stage expressing tagged cuticle collagen SQT-3::mNG over 200 seconds. Scale bars, 20 µm (xy of **a, e, g**), 5 µm (xz of **a**), 10 µm (**b**).

## Discussion

In this manuscript, we present a novel inline OPM design that eliminates the need for a separate inclined tertiary microscope, as the secondary microscope effectively serves the tertiary microscope’s function. This design significantly simplifies the OPM system and corrects aberrations caused by tilts. Our RIDE-OPM design incorporates a mechanical focuser with a small mirror attached: a departure from traditional OPM setups (**Supplementary Fig. 16**). By carefully selecting the step size for the mirror movement and an appropriate focuser travel range, RIDE-OPM can image the desired z-depth effectively. We found that for a small travel range of the focuser to cover our FOV, RIDE-OPM can operate at the camera-limited speed of our system. This design is versatile, and adaptable to any OPM configuration involving various primary and secondary objective pairs, with the selection of tube lenses for optimal pupil matching being the only major consideration. This creates a more compact system and reduces major drift sources seen in traditional OPM. While minor drift in the secondary objective may occur, the reflective mirror compensates for the focus drift allowing for long-term, unsupervised imaging (**Supplementary Movie 5**). Its inline optical design also avoids tilt-induced aberrations, ensuring stable, high-quality imaging over long durations (**Supplementary Fig. 17**). While previous inline, reflection-based designs have attempted to eliminate the tertiary objective, they have typically been limited by trade-offs in numerical aperture (NA), volumetric imaging capability, or imaging speed. Although mechanically integrated interfaces between O_II_ and O_III_ can improve stability in conventional OPM systems, such approaches are not easily adaptable to varying primary objective NAs and tilt angles simultaneously^45^.

In the current implementation, the RIDE-OPM design incurs an approximately 50% loss of collected fluorescence due to the use of an optical isolator. As a result, the system may exhibit a reduced signal-to-noise ratio (SNR) compared to conventional OPM configurations. Future developments aimed at minimizing or eliminating this loss, including approaches for near lossless fluorescence collection, are expected to further improve the SNR performance of the system. With that said, even at its current design, the efficiency of light preservation at the O_II_ - O_III_ interface still makes RIDE-OPM effective for biological imaging. Our results show that the fluorescent signal captured using our setup is strong enough to reliably image a wide range of specimens with the sCMOS camera we employed, without requiring signal amplification or specialized detection schemes. Another issue with this implementation is the generation of out-of-focus light from the features in neighbouring planes of the remotely focused plane. Although RIDE-OPM generates some out-of-focus light, its impact is less significant compared to the wide-field systems. This reduced impact is due to the system’s output PSF being a product of both the illumination and detection PSFs^46^ (**Supplementary Fig. 18**). The optical sectioning capability provided by the LS in OPM results in a sharper PSF compared to those found in wide-field detection systems^40^. Looking ahead, we anticipate that future designs employing innovative techniques will capture all emitted fluorescence and eliminate out-of-focus light, enhancing optical sectioning. In summary, OPM has garnered interest within the imaging community due to its advantages, such as open-top sample mounting and the capability for fast, high-resolution imaging. However, challenges like complex geometries, the need for sophisticated and costly objectives, tilt-dependent performance, and potential NA loss have driven researchers to seek simpler inline solutions. Our RIDE-OPM system addresses these issues by offering a straightforward optical setup that utilizes readily available off-the-shelf components, making it easily integrable with any OPM microscope. We are confident that RIDE-OPM will prove to be a valuable asset for any biomedical imaging facility requiring rapid and high-resolution capabilities.

## Methods

### Optical setup

RIDE-OPM was constructed on an optical table using an open-top sample mounting geometry (**Supplementary Fig. 19, 20**). The system incorporated four collinear but independently modulated laser channels at wavelengths of 405 nm, 488 nm, 561 nm, and 637 nm (LX 405, LX 488-50C, LS 561-50, and LX 637-140C, Coherent OBIS). These laser beams were combined within a software-controlled integrated laser box, allowing for future expansion. The beams were merged using three dichroic mirrors, cleaned, and collimated by a 50 µm pinhole (P50D, Thorlabs) and a pair of 50 mm achromatic lenses.

To create a Gaussian LS, the collimated beam was expanded eightfold in one dimension using a pair of cylindrical lenses and then focused in the orthogonal dimension using a third cylindrical lens before reaching a resonant galvanometric mirror. The resonant galvo, conjugated to the sample plane, oscillated the LS along the x-axis to minimize shadowing effects caused by light obstruction. We placed an adjustable mechanical slit conjugated to the back pupil of the illumination objective, controlling its effective NA. The transmitted light after the slit was imaged by a galvanometric mirror (GM) which translates the LS along y-axis for 3D image acquisition. To maintain plane conjugation, GM shared a common one-dimensional translation stage with another mirror. A one-dimensional mechanical translation stage was used to shift the optical axis by a few millimeters from the inline optical axis, ensuring that the beam hit off-center in the illumination objective and thereby creating a tilted LS in the sample space.

In this implementation, the oil immersion objective was chosen due to its widespread adoption and proven performance over the past two decades in both high-resolution and super-resolution imaging^8,41,47,48^. However, it is important to note that the RIDE-OPM technique is not restricted to oil immersion objectives and can be adapted for use with other types as well. Specifically, we used a Nikon Plan Apo 60×/1.4 (scale 1) and 40×/1.0 (scale 2) oil immersion objective as our primary objective (O_I_), which also served to collect the fluorescence emitted from the specimen. For direct imaging, a dichroic mirror, mounted on a flip stage, separated the fluorescence signal from the excitation light and directed it toward the camera. This setup allowed us to visualize 1D lines within the confocal range of the LS by shifting the specimen along the z-axis, causing these lines to shift by a few pixels.

For the RIDE-OPM set up, a quad-band filter was placed in the optical path to separate the excitation and fluorescence light. To capture the entire xz’ sample plane, we implemented a remote focusing arrangement between O_I_ and O_II_ with perfectly matched pupil lenses (L5-L10). The secondary objective (O_II_) was Nikon 40×/0.95 NA (scale 1) and 60×/0.7 NA (scale 2) air objective, which ensures the entire aperture angle of O_I_ gets captured. This remote focusing geometry created a virtual image of the xz’ plane with identical lateral and axial magnification (1.5×), following the refractive index ratio between the two media^31^.

The scanning illumination beam was descanned for fluorescence emission using the same GM, ensuring that all xz’ planes remained stationary along the y-axis after passing through O_II_. To reflect light from different z-planes within the virtual image created by remote focusing, we employed a high-speed linear focuser.

A polarizing beam splitter (PBS) was used to separate the S- and P-polarized components of the unpolarized fluorescence light. In scale 1, the P-polarized light passed through a quarter-wave plate (QWP), converting it into circularly polarized light before entering O_II_. This light then formed the virtual image of the tilted xz’ plane. A mirror, placed orthogonally to the optic axis, reflected this light back, causing it to undergo another phase shift via the QWP, transforming it into S-polarized light. The PBS then reflected this S-polarized light, which was subsequently focused onto the camera by a tube lens (TL)/achromatic lens (L11), forming a 1D line of the 2D xz’ plane. We chose S-polarized light for imaging in scale 2, confirming that imaging performance were preserved.

To capture the entire xz’ sample plane, we dithered the mirror at an amplitude covering the virtual image with Nyquist sampled step at a speed at least twice that of the camera’s frame acquisition rate (**Supplementary Note 6**). This allowed us to image the entire 2D xz’ sample plane using a scientific complementary metal-oxide semiconductor (sCMOS) camera. For multi-color imaging, a high-speed optical filter wheel was installed before the camera, equipped with four emission filters for violet, green, red, and far-red channels. A sequential and synchronous modulation of the laser lines and the emission filters is used to image multiple channels stained with different fluorophores. A tube lens (TL)/achromatic lens (L11) was used to achieve a final magnification of 60× (scale 1) and 41.2× (scale 2), resulting in a pixel size of 108 nm and 160 nm, respectively. A 3D motorized stage was used to translate the sample in all three dimensions for multi-position imaging. The specimen was placed on a 1.5 coverslip with a thickness of 0.17 mm.

The timing diagram for the microscope equipment is shown in **Supplementary Fig. 21**. A detailed comparison of mirror-based reflection detection methods is provided in **Supplementary Table 1**. Optical schematic of the complete RIDE-OPM system and a comprehensive list of components used in building the RIDE-OPM system are shown in **Supplementary Fig. 22** and **Supplementary Table 2**, respectively.

### Photobleaching comparison among existing OPM, RIDE-OPM, confocal and ASLM system

The existing OPM system was custom-built in-house, following the optical design detailed by Sapoznik et al.^13^. The system utilized a Nikon Plan Apo 100×/1.35 NA silicone oil immersion objective as the primary, a Nikon S Fluor 40×/0.9 NA air objective as the secondary, and a Calico AMS-AGY v2.0 as the tertiary objective. This configuration achieved a resolution comparable to that of RIDE-OPM scale 1.

Additionally, we constructed an ASLM system in-house, based on the schematic presented by Chakraborty et al^49^. For both illumination and detection, we employed an ASI multi-immersion objective (0.7 NA at refractive index 1.45), enabling symmetric and high-resolution imaging.

We also used commercially available confocal microscope (Zeiss LSM 980). To fair comparison we use 63×/1.4 NA objective to achieve a similar resolution of RIDE-OPM scale 1.

### Microscope control and data acquisition

The microscope system was controlled and images were acquired using a Windows-operated Dell Precision 7920 computer. This system is powered by two Intel® Xeon® Silver 4210R CPUs, running at 2.40 GHz and 2.39 GHz, and is equipped with 128 GB of memory for processing microscopic data. It operates on a 64-bit architecture, with two sockets supporting 20 logical processors. For graphical processing, the system includes an NVIDIA Quadro RTX 4000 GPU, featuring 8 GB of dedicated memory and 63.8 GB of shared memory, resulting in a total GPU memory of 71.71 GB. The microscope control software is based on LabVIEW 2020 64-bit, incorporating the Vision Development Module and LabVIEW. The software interfaced with the camera (Flash 4.0, Hamamatsu) through the DCAM-API for an active Silicon Firebird frame grabber and generated a series of deterministic transistor–transistor logic (TTL) triggers via a field-programmable gate array (PCIe-7852R, National Instruments). This synchronized control ensured precise timing between image acquisition, illumination, and stage movement. Subvolume acquisition was performed by translating the specimen using a three-axis motorized stage (3DMS and MP-285A, Sutter Instruments) for fine and rapid volumetric positioning.

### Sample preparation

#### (a) 100 and 500 nm fluorescent beads

10 ml fluorescent bead (F8803 for 100nm and F8813 for 500 nm, invitrogen) stock solution was prepared at 1:10 dilution from the manufacturer stock. From this stock, 770 µl bead dilution at 1:10 was transferred in a centrifuge tube and sonicated for an hour. Next, 50 µl of the sonicated bead dilution was transferred to a new centrifuge tube and vortexed for 3 minutes. 1 ml of glycerol (GX0185-6, Sigma Aldrich) was then added, followed by 5 minutes of vortexing. The solution was left to incubate overnight at room temperature. 1/2 drop (∼20 µl) of the final solution was placed onto a 1.5 coverslip (0.17 mm thick) for imaging.

#### (b) *C. elegans* culture conditions and sample preparation

*C. elegans* were maintained on Nematode Growth Media with a standard diet of *E. coli* OP50 until they reached the target developmental stage. Synchronized animal populations were obtained by isolating embryos through sodium hypochlorite treatment of gravid adults, followed by incubation in M9 buffer for 20 hours to allow hatching. The *C. elegans* strains used in this study are listed in **Supplementary Table 3**. For NucSpot staining, animals were fixed with 100% methanol for 20 minutes. After fixation, the methanol was removed, and the animals were permeabilized with 0.2% Triton X-100 for 30 minutes. Worms were then washed once with PBS. NucSpot 650/665 (Biotium) was added at a 1:1000 dilution in PBS and incubated for 20 minutes.

For imaging, *C. elegans* were mounted on 3% agarose pads in M9 buffer. When necessary, specimens were immobilized using 30 mM sodium azide.

#### (c) Adult Drosophila brain

Adult flies were dissected in ice-cold Schneider’s insect media (Sigma Aldrich). Dissected brains were then fixed in 4% paraformaldehyde (PFA, EMS) prepared in 1X PBST (phosphate-buffered saline with 0.5% Triton X-100) for 27 minutes at room temperature. After fixation, brains were washed three times, each for 20 minutes, in 1X PBST. They were then incubated in a blocking solution made of PBST with 2.5% Normal Goat Serum and 2.5% Normal Donkey Serum (Jackson ImmunoResearch) for 40 minutes at room temperature. Brains were incubated with primary antibodies at 4°C —for two nights. After primary antibody incubation, brains were washed three times (20 minutes each) in PBST and then incubated with secondary antibodies for two nights at 4°C. Following secondary antibody incubation, brains were again washed three times in PBST (20 minutes each). After washing, brains were mounted using DPX mounting medium, following the Janelia FlyLight protocol. Chicken anti-GFP was diluted at a ratio of 1:1500, and mouse anti-Bruchpilot (nc82) was diluted at a ratio of 1:50.

#### (d) Cell lines and cell staining

All cell lines were maintained under standard culture conditions (37°C, 5% CO_2_) using their specific growth media: MDA-MB-231 and EO771 cells were cultured in RPMI-1640 and DMEM medium containing 10% fetal bovine serum (FBS) and 1% penicillin/streptomycin, respectively. MCF10A cells were grown in DMEM/F12 medium supplemented with 5% horse serum, 20 ng/mL EGF, 0.5 μg/mL hydrocortisone, 10 μg/mL insulin, and 1% penicillin/streptomycin. For immunofluorescence experiments, cells were seeded in 4-well chamber slides at 30-40% confluency and allowed to attach overnight. After 24 hours, cells were fixed with 4% paraformaldehyde for 15 minutes at room temperature. Following fixation, cells were blocked with 3% bovine serum albumin (BSA) containing 0.04% saponin for 20 minutes. Primary antibodies against GLUT3 (anti-mouse) and β-Tubulin (anti-rabbit) were diluted 1:200 in blocking solution (3% BSA with 0.04% saponin) and applied to the cells overnight at 4°C. The next day, the primary antibody solution was removed, and cells were washed three times with PBS (5 minutes per wash). Secondary antibodies (Alexa Fluor 488 anti-rabbit and Alexa Fluor 561 anti-mouse) were then applied for 1 hour at room temperature, followed by three additional PBS washes.

For F-actin visualization, cells were stained with phalloidin conjugated to Alexa Fluor 647 (400X) for 30 minutes. Finally, cells were mounted with DAPI-containing mounting medium and prepared for microscopic analysis.

#### (e) Cloning, media and cell growth of *E. coli*

Standard molecular biology techniques were used to construct pIB184_Km_sfGFP. The gene encoding superfolder green fluorescent protein (sfGFP) was amplified from a pET DUET_sfGFP plasmid by polymerase chain reaction using forward (TGATGTTGGATCCCCGCAGGAGATATACAAATGAGCAAAGGAGAAGAACTTTTC) and reverse (TATCGATAGATCTCGAGCTCTAGTTATTTGTAGAGCTCATCCATGCCATG) primers and Q5 high-fidelity DNA polymerase. The amplicon was inserted into the SacII and EcoRI restriction sites of pIB184_Km^50^ with isothermal assembly and transformed into *E. coli* DH10B-T1^R^ chemically competent cells. The identity of the cloned insert was verified by sanger sequencing. pIB184-Km was a gift from Indranil Biswas (Addgene plasmid # 90195 ; http://n2t.net/addgene:90195 ; RRID:Addgene_90195).

To express sfGFP, pIB184_Km_sfGFP was transformed into *E. coli* BL21(de3)-T1^R^ chemically competent cells. A single colony was used to inoculate a 10 mL overnight culture of LB media with 50 µg/mL kanamycin at 37°C with shaking at 200 rpm. Due to presence of P23 constitutive promotor sfGFP was expressed and was visible in the cell. Cells were collected from 1 ml of the overnight culture by centrifugation at 16000 × g for 1 min at room temperature. The cells were washed with phosphate buffered saline (PBS) and centrifuged again. 20 μl of cell suspension was diluted with 980 μl of PBS. 4 μl of this cell suspension was spread on the cover slip for observation on the microscope.

### Image processing pipeline

The multi-dimensional image stacks were saved in .tif format to a shared drive connected via 10-gigabit Ethernet. As each 3D stack is acquired at an angle relative to the xy-plane, images were deskewed using Pyclesperanto, a Python-based image processing library^51^. Deconvolution was performed using a custom MATLAB implementation of the standard Richardson-Lucy algorithm^52,53^. We used fiji^54^, an open-source software package for visualizing images. The Bigstitcher^55^ was used to stitch the image tiles. *E. coli* were segmented using StarDist^56^, and their motion was tracked with TrackMate^57^.

## Data availability

The report and its Supplemental Information provide the primary results that underpin the findings of this study. The dataset used for this study is available upon request to the corresponding author.

## Acknowledgements

This project was supported by the National Institutes of Health (Grant No. R35GM151152 to T.C, NINDS R01NS136555 to M.H.S., and R35GM142530 to M.C.W.). Additional support was provided by the NSF career award (IOS-2047020) to M.H.S., the National Cancer Institutes and the Cancer Center Support grant (P30CA118100) to B.C.M., and the American Cancer Society (ACS) – Institutional Research Grants (IRG) to T-H.K. We also acknowledge funding from the University of New Mexico Pathology Start-up grant.

## Author contributions

M.N.H.P. and T.C. designed the research. M.N.H.P. built and operated the microscope. M.N.H.P., W.H., and N.S. performed experiments and image analysis. N.S. built ASLM system and C.S. built the existing OPM system. M.N.H.P. and B.C.M. imaged the *C. elegans*. B.C.M. and O.P. prepared the *C. elegans* specimen, A.Q.B. and T-H.K. prepared multi-stained cell lines, A.R.W. and M.H.S. prepared Drosophila brain samples, and M.R.I.R. and M.C.W. prepared *E. coli* specimen; and guided for imaging of those specimens. M.N.H.P. and T.C. wrote the manuscript. All authors read and provided feedback and necessary input on the final manuscript.

## The authors declare the following competing interests

T.C. and M.N.H.P. have filed a patent application (United States Patent and Trademark Office application) for the RIDE-OPM microscope mentioned here. The remaining authors declare no competing interests.

