## Supplementary Information for "Reflected Inline Detection in Epi Oblique Plane Microscopy"

**Supplementary Note 1. Positioning of RIDE-OPM relative to the static mirror-based approach.** Static mirror–based inline detection approaches, such as single-objective oblique plane microscopy (obSTORM) and dual-view oblique plane microscopy (dOPM), employ a mirror positioned at an angle relative to the optical axis after the secondary objective. This angular configuration inherently causes the fluorescence emission to deviate from the optical axis, leading to a loss of collected signal and a reduction in effective numerical aperture (NA).

In contrast, RIDE-OPM utilizes a dithering mirror positioned orthogonally to the optical axis, enabling reflection of the full fluorescence signal while preserving its propagation direction. This design maintains the full NA across all one-dimensional lines along the axial extent of the tilted virtual image plane, allowing the complete two-dimensional tilted plane to be captured without compromising imaging performance.

Additionally, obSTORM is limited to imaging a single tilted 2D plane and does not support volumetric imaging. Its reduced NA further limits spatial resolution; although super-resolution can be achieved via dSTORM, this relies primarily on fluorophore blinking and reconstruction algorithms rather than the optical design itself. Similarly, dOPM employs a motorized translation stage with a custom prism–mirror assembly, increasing system complexity, reducing imaging speed, and introducing additional computational steps for image fusion. The requirement for precise mechanical motion also introduces potential instability.

In contrast, RIDE-OPM employs dithering of the fluorescence signal to reconstruct the tilted imaging plane without sacrificing resolution, speed, or system stability. Importantly, this dithering mechanism does not introduce performance trade-offs. A piezoelectric mirror is used to both scan the light-sheet and de-scan the emitted fluorescence, enabling volumetric imaging over the desired axial range. A quantitative comparison of reflection-based inline imaging approaches, highlighting the performance advantages of RIDE-OPM, is provided in Supplementary Table 1.

**Supplementary Note 2. Relative collection efficiency (RCE) calculation for RIDE-OPM.** The red cone represents the collection cone of the secondary objective (OII; NA = 0.95, air), corresponding to a half-angle of ~71.8°, while the blue cone represents the collection cone of the primary objective (OI; NA = 1.4, oil immersion), corresponding to a half-angle of ~67.5°. The overlapping region, shown in yellow, defines the effective collection solid angle of the system ($\Omega_{OPM}$) and is used to quantify the relative geometric collection efficiency. $\Omega_{OPM}$, was determined from the overlap of the objective acceptance cones on the unit sphere; for the present RIDE-OPM geometry, both $\Omega_{OPM}$ and the collection solid angle of the primary objective alone, $\Omega_{O1}$, correspond to a half-angle of 67.5°. Optical transmission losses were incorporated through the relay optics transmission and the polarization-dependent transmission of the PBS/QWP isolator. The relay transmission was calculated using $T_{O2/O3} = 0.9$, and the PBS transmission factor was taken as $T_{PBS} = 0.5$, corresponding to a 50% loss of fluorescence due to polarization selection. Thus, the RCE was evaluated relative to the primary objective collection solid angle while accounting for both geometric overlap and downstream optical transmission losses.

**Measurement of the relative collection efficiency of RIDE-OPM. Supplementary Fig. 4** represents the collection solid angles mapped onto the surface of the unit sphere. The red cone represents the collection cone of the secondary objective ($O_2$; NA = 0.95, air), corresponding to a half-angle of ~71.8°, while the blue cone represents the collection cone of the primary objective ($O_1$; NA = 1.4, oil immersion), corresponding to a half-angle of ~67.5°. Both cones are projected into a common coordinate system aligned with the optical axis (black line). The overlapping region, shown in yellow, defines the effective collection solid angle of the system ($\Omega_{OPM}$) and is used to quantify the relative geometric collection efficiency.

The geometrical collection efficiency is determined numerically by calculating the area of intersection between the acceptance cones of the objectives on the unit sphere. In conventional dual-view OPM (dOPM), this overlap includes contributions from three objectives ($O_1$, $O_2$, and $O_3$), whereas in the RIDE-OPM architecture the reflective design eliminates the tertiary objective, resulting in a two-objective system in which fluorescence collected by $O_{II}$ is reflected and re-imaged through the same optical pathway. This configuration preserves the full angular acceptance of the system, enabling tilt-independent NA utilization and effectively eliminating geometric losses associated with tertiary objective interfaces.

In addition to geometric considerations, the overall collection efficiency incorporates transmission losses in the detection path. These include polarization-dependent losses introduced by the polarizing beam splitter (PBS) and quarter-wave plate (QWP), which are accounted for through a transmission factor that depends on the polarization characteristics of the emitted fluorescence (e.g., ~50% transmission for rapidly tumbling fluorophores and higher values for partially polarized emission). The efficiency also accounts for transmission through the relay optics, including the double pass through the secondary objective ($O_{II}$) and experimentally measured transmission of the PBS–QWP assembly.

The final relative collection efficiency is therefore defined with respect to the collection solid angle of the primary objective alone ($\Omega_{O1}$), incorporating both the geometrical overlap of the cones and the cumulative transmission efficiency of the optical system. This framework enables a direct comparison between RIDE-OPM and conventional OPM configurations, highlighting the improved geometric efficiency and tilt-independent performance of the reflective design.

**Supplementary Note 3. Axial dithering to image a 2D plane.** The core concept of RIDE-OPM is based on transforming the axial dimension into a lateral dimension by optically encoding the depth information into a tilted 2D virtual image plane. This tilted plane, which contains oblique axial information, is projected onto the camera sensor within a single camera exposure, allowing for volumetric imaging without mechanical z-scanning.

Under ideal conditions, a static mirror placed orthogonally to the optic axis in the remote detection path (after $O_{II}$) can reflect light from a specific axial section of the tilted virtual image. However, due to the intersection of two non-parallel planes, the mirror plane and the virtual image plane, only a 1D line, the line of intersection between the two planes, will satisfy the conditions for proper reflection. Fluorescence emitted from this line is redirected back through the optical system and ultimately imaged onto the sCMOS camera, resulting in the formation of a partial image corresponding to that depth.

By adjusting the axial position of the mirror, different 1D lines across the virtual tilted plane can be selectively reflected and recorded. This principle enables stepwise acquisition of the entire 2D image plane through sequential 1D line reflections. However, to achieve real-time capture of the full tilted image within a single exposure time, we implemented rapid axial dithering of the mirror, oscillating it back and forth at a frequency at least twice the rate of the camera exposure time. This ensures that the mirror traverses the entire depth of the light sheet's (LS's) confocal range within each exposure cycle. The net effect is a time-averaged integration of all 1D slices, forming a complete and continuous image of the tilted plane (**Fig. 1e**).

To enable efficient and isolated detection of the reflected fluorescence, we employed an optical isolator in the detection path. This isolator consisted of a polarizing beam splitter (PBS) and a quarter-wave plate (QWP). The PBS ensures that only a specific polarization of light passes through, while the QWP rotates the polarization of the reflected light by 90° (via twice phase rotation), allowing it to exit the system through a different path and be directed to the camera, effectively separating incoming and outgoing optical signals.

The effective point spread function (PSF) of the system is the product of the illumination PSF and the detection PSF (**Supplementary Fig. 6**). This system PSF governs the resolution limits and is central to defining the optimal strategy for dithering.

The dithering step size, which dictates how finely the mirror moves during axial oscillation, is defined by the system's PSF. To faithfully reconstruct spatial features without aliasing, the step size must comply with the Nyquist sampling criterion (**Supplementary Fig. 5**). By adhering to this criterion, RIDE-OPM enables accurate spatial representation of the obliquely imaged volume while maintaining the temporal advantages of single-exposure acquisition.

**Supplementary Note 4. Potential drift correction in RIDE-OPM.** OPM systems are designed for high-resolution imaging at camera-limited acquisition speeds. While state-of-the-art OPM systems are capable of delivering high spatial resolution and camera-limited imaging speeds, their long-term performance is compromised by time-dependent drift. This drift arises primarily from temperature-induced expansion, mechanical vibrations, and environmental instabilities that affect the precise optical alignment required for the system to function optimally[1,2]. The most critical alignment constraint in conventional OPM involves maintaining the intersection of the tilted virtual image plane, formed after $O_{II}$, with the NFP of $O_{III}$ (**Supplementary Fig. 3a**). Even a minor axial displacement of the NFP of the tertiary objective can lead to misalignment between the image plane and the detection focal plane, introducing significant defocus and broadening of the PSF (**Supplementary Figs. 3b–3c**). This issue becomes especially problematic in systems that use high NA objectives, where the depth of field is extremely shallow, on the order of hundreds of nanometers. Under these conditions, submicron drift is sufficient to degrade resolution and image fidelity, severely limiting the system's ability to support extended imaging sessions required for live or developmental studies.

The predominant source of this drift is the tertiary objective assembly itself, which is susceptible to both axial and lateral displacement over time. In an effort to mitigate this, some implementations have introduced active drift correction mechanisms, such as mounting the tertiary objective on a piezoelectric actuator to continuously realign the focus during imaging[2,3]. While this approach can reduce the impact of axial drift, it adds substantial mechanical and control complexity to the system. Furthermore, the dynamic correction range is often limited, and the added components can reduce the system's temporal responsiveness and robustness—especially problematic when imaging time-sensitive biological processes. As a result, the reliable imaging performance of conventional OPMs is typically limited. Additional instability may arise from lateral drift of the virtual LS image formed after the secondary objective, further contributing to PSF degradation.

The RIDE-OPM system resolves these limitations through a fundamental reconfiguration of the detection path. By eliminating the tertiary objective entirely, RIDE-OPM removes the primary mechanical source of axial drift. Instead of imaging through a tilted tertiary detection arm, RIDE-OPM uses an orthogonally oriented mirror to reflect the tilted virtual image back toward the camera. This approach preserves a common optical axis throughout the system (**Supplementary Fig. 3d**), ensuring geometric stability and avoiding the alignment sensitivity associated with a tertiary objective's NFP.

Moreover, the compact and symmetric optical architecture of RIDE-OPM allows the system to passively resist drift-induced misalignment. Even minor lateral shifts of the virtual image due to movement in the secondary objective can be absorbed without significantly affecting the system's performance. As a result, RIDE-OPM provides a highly stable and drift-tolerant imaging platform, ideal for long-term, unsupervised imaging applications such as time-lapse developmental biology.

In summary, the streamlined optical configuration of RIDE-OPM not only simplifies system construction and alignment but also delivers superior mechanical robustness over time, making it well-suited for continuous high-resolution imaging across extended durations without requiring active drift correction or frequent recalibration.

**Supplementary Note 5. Implementation of various oblique-angle illuminations.** In OPM, the properties of the light-sheet (LS) inherently create trade-offs between axial resolution and imaging depth. This is primarily due to the inverse relationship between the LS thickness and its confocal range. Additionally, increasing the LS tilt angle, which is often necessary to access greater imaging depths above the coverslip, results in numerical aperture (NA) loss in conventional OPM systems, further compromising lateral resolution. As a result, OPM systems must balance three competing factors: LS tilt angle, axial resolution, and imaging depth.

Most existing OPM systems operate at a fixed LS tilt angle, typically between 30° and 45°, to manage this trade-off. However, once the LS tilt is changed, the corresponding virtual image formed after the secondary objective ($O_{II}$) also shifts in tilt. Since the tertiary objective ($O_{III}$) must have its narrow nominal focal plane (NFP) aligned with this tilted virtual image for optimal imaging, adjusting the tilt requires a complex and precise repositioning of the entire tertiary imaging arm ($O_{III}$, tube lens, and camera). This includes not just rotational adjustment but also two-dimensional linear translation, which is technically challenging and often impractical (**Fig. 5a**).

RIDE-OPM overcomes these limitations by eliminating the need for a tertiary objective. It maintains consistent resolution near the NFP of $O_I$ across a range of LS tilt angles, preserving both fluorescence signal and effective NA even as the tilt varies (**Fig. 1g** and **Supplementary Fig. 2**). Switching the LS tilt angle in RIDE-OPM is achieved simply by translating Linear Stage 1 (LST 1) (**Fig. 5b**) and adjusting the axial dithering range of the mirror. No camera or objective repositioning is needed.

This simplicity allows even non-expert users to change the LS tilt angle, and thus the imaging depth, via software, making RIDE-OPM a versatile and user-friendly system. However, each tilt angle does require adjustment in the deskewing parameters during image processing, which will result in different z-depths and output image sizes.

In this work, we focused on validating RIDE-OPM's performance within the conventional 30°–45° tilt range, demonstrating its ability to maintain both fluorescence signal and NA throughout. While RIDE-OPM achieves spatial resolutions of approximately ~318 nm (x) and ~300 nm (y) for both tilt angles, the existing OPM system, even at a 30° light-sheet tilt, achieves resolutions of ~300 nm (x) and ~336 nm (y)[1]. Future studies could explore even steeper oblique angles. For example, Botcherby et al. showed that imaging the meridional plane (i.e., at a 90° tilt) requires an additional scanning element for proper image formation[4]. RIDE-OPM may offer a pathway toward such advanced configurations with further development.

**Supplementary Note 6. Simultaneous operation of the resonant and the dithering mirror.** In the RIDE-OPM system, axial dithering mechanism is employed to enable high-resolution, time-integrated imaging without performance interference. The dithering mirror is responsible for reflecting the fluorescence signal back toward the detection path. It oscillates axially at a rate at least twice as fast as the camera exposure time, effectively enabling time-integrated acquisition of the tilted virtual image plane during a single frame exposure. This mechanism is critical for capturing a complete 2D projection of the inclined focal plane without requiring multiple exposures.

In parallel, a resonant mirror (CRS, Cambridge Technology) is placed at a plane conjugate to the sample and is used to laterally scan the LS across the specimen (xz'). This mirror operates at a frequency of several kilohertz, an order of magnitude faster than the dithering mirror, to ensure uniform LS illumination and minimize striping artifacts.

Because the two mirrors operate at distinct spatial axes (axial for dithering, lateral for resonant scanning) and widely separated temporal frequencies, they function independently and do not interfere with each other's operation. This decoupling ensures that both mechanisms can perform their intended roles, dithering for axial integration and resonant scanning for LS uniformity, reliably and simultaneously.

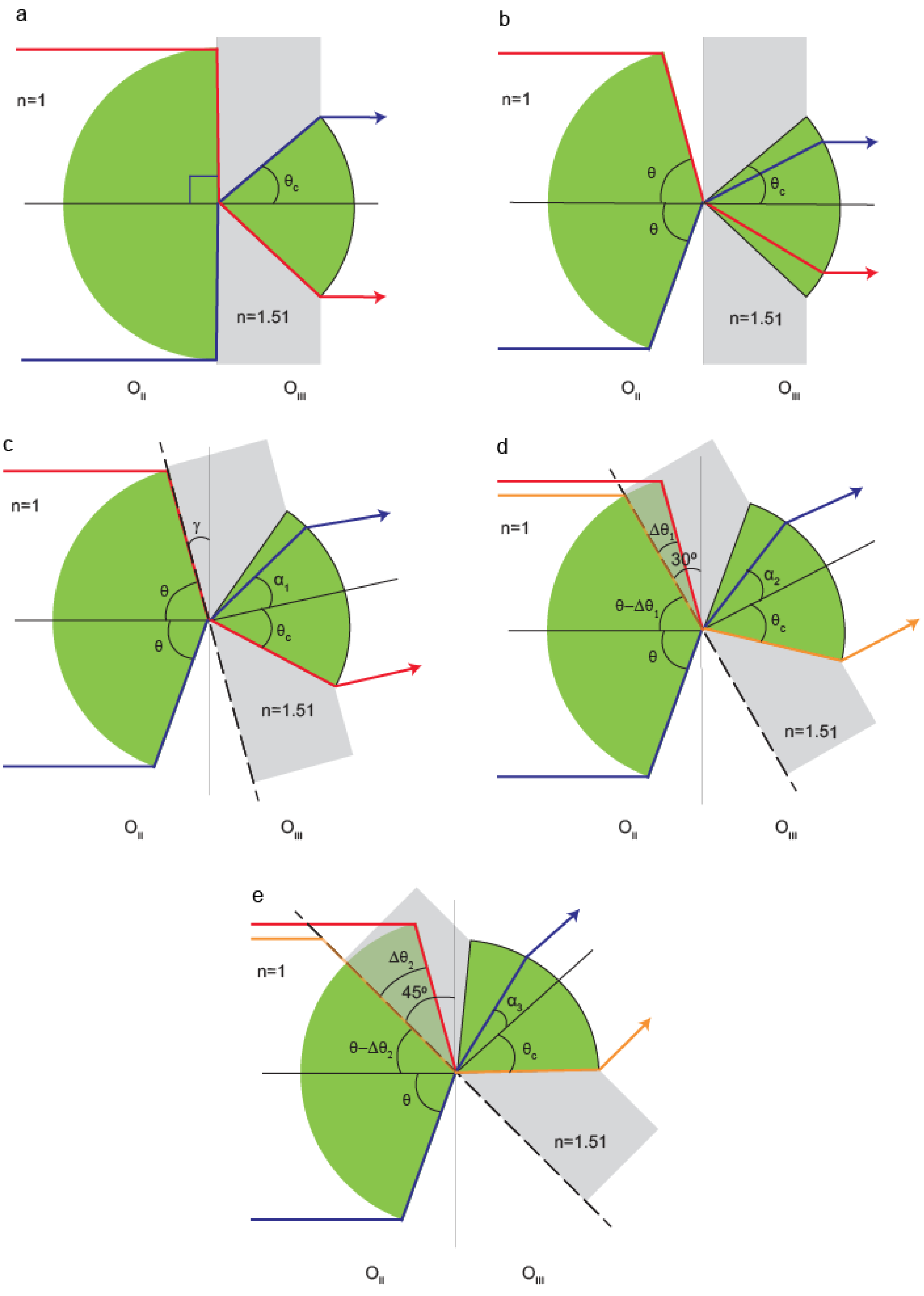


$\alpha_1 > \alpha_2 > \alpha_3$

**Supplementary Fig. 1 | Tilt dependent NA loss at the $O_{II}$ - $O_{III}$ interface for 1 NA $O_{III}$ (Snouty) objective.** The green area represents the light cone at the $O_{II}$ - $O_{III}$ interface, while the gray area signifies the glass piece (RI 1.51) positioned in front of $O_{III}$. Red, blue, and yellow are used to depict different light rays. **a-b,** In-line placement of $O_{II}$ and $O_{III}$ (0° tilt of $O_{III}$). Light coming from $O_{II}$ at 90° of half aperture angle results in entering $O_{III}$ at the critical angle ($\theta_c$) due to the glass interface of $O_{III}$ (**a**) while a Lower incoming angle leads to entry below $\Theta_c$ (**b**). **c-e,** Placement of $O_{III}$ at tilt angles γ° (**c**), 30° (**d**) and 45° (**e**). At γ° of tilt, the red light ray enters $O_{III}$ at $\theta_c$ angle while the blue light enters at $\alpha_1$ angle ($\alpha_1 < \theta_c$). At 30° tilt, yellow light ray enters at $\theta_c$, and blue light enters at $\alpha_2$ ($\theta_c > \alpha_1 > \alpha_2$), showing a decline in effective NA represented by the half aperture angle of $\Delta\Theta_1$. This loss of effective NA is more pronounced at 45° tilt, indicated by an even greater loss of half aperture angle ($\Delta\Theta_2$), as angles progressively decrease from θc through α1, α2 to α3, highlighting the increased NA loss with greater tilt angles.

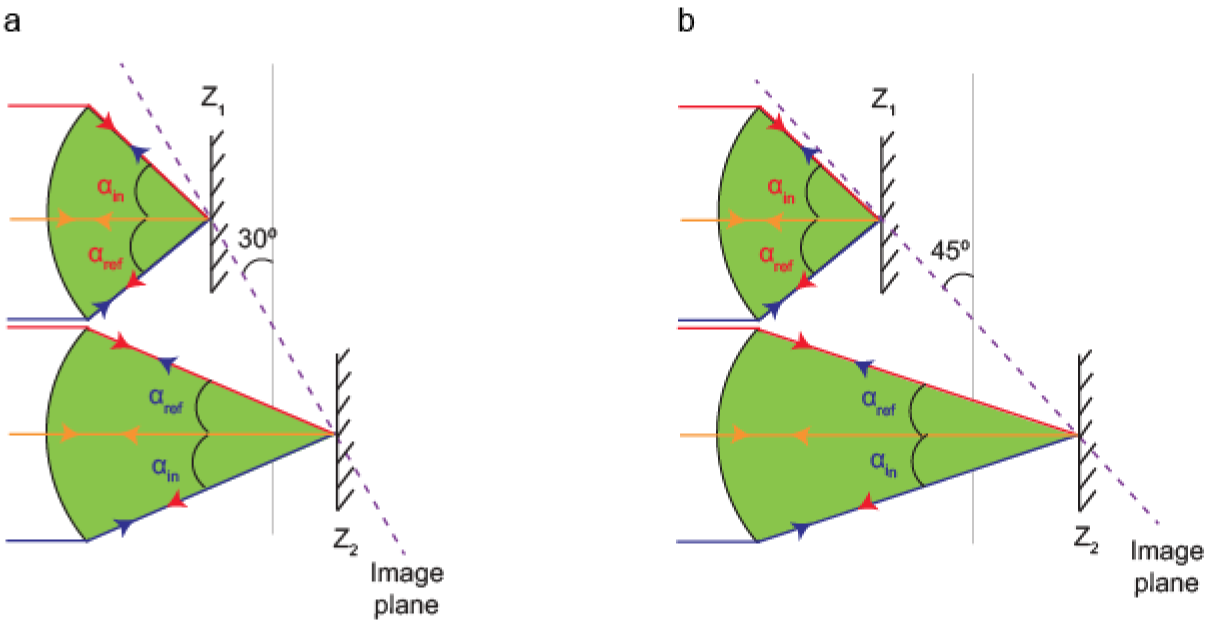


**Supplementary Fig. 2 | Tilt independent light path in $O_{II}$ for RIDE-OPM. a-b,** Propagation of fluorescence light at $O_{II}$ - $O_{III}$ interface is presented for LS tilts of 30° (**a**) and 45° (**b**). The green area depicts the incoming and reflected light cone at OII. The red, blue, and yellow lines represent two marginal rays and the principal ray of the cone, respectively. In both scenarios, the incoming and reflected light paths preserve the same half aperture angle, ensuring consistent imaging without light loss despite the tilt adjustments.

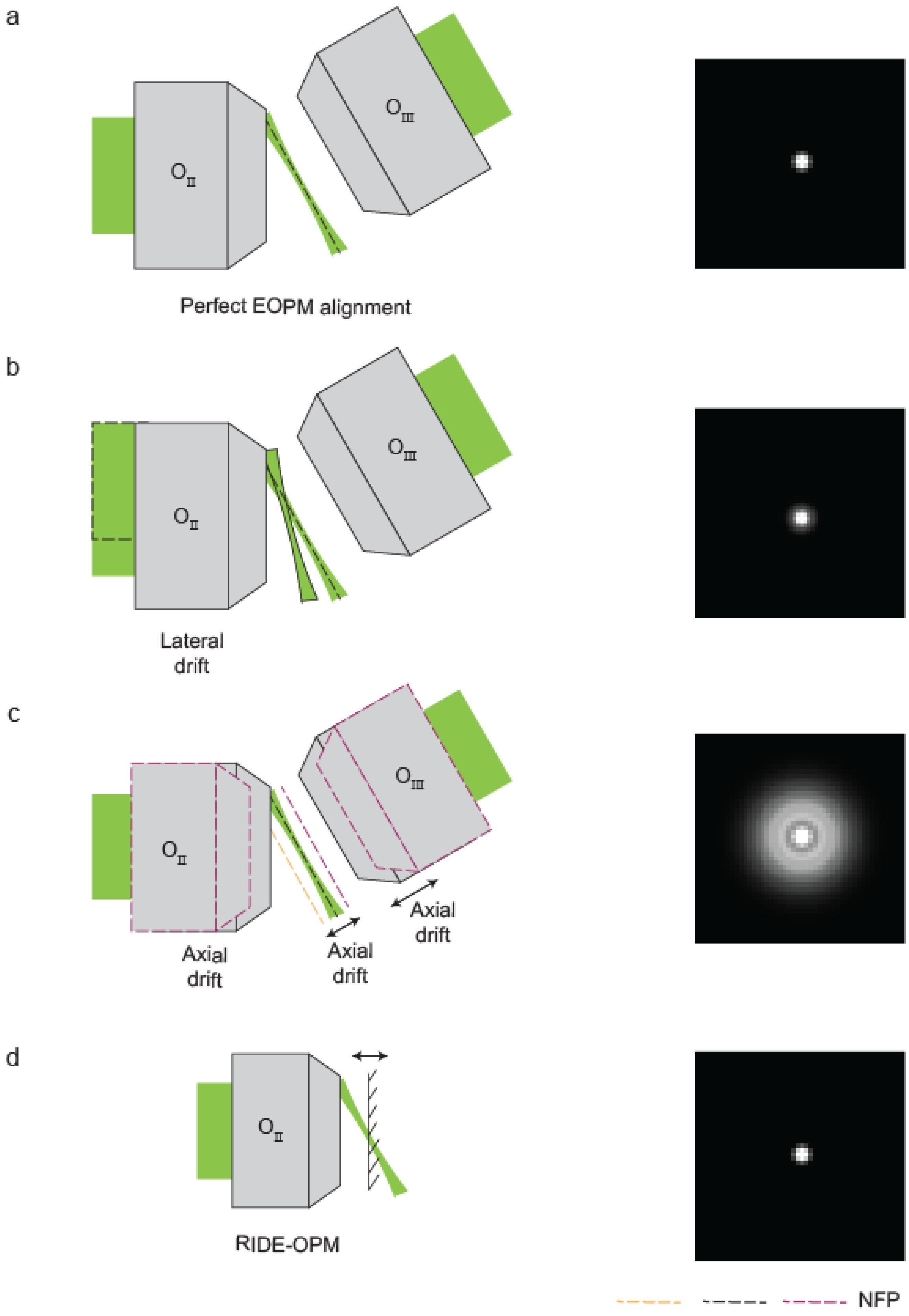
a
$O_{II}$
$O_{III}$
Perfect EOPM alignment
b
$O_{II}$
$O_{III}$
Lateral drift
c
$O_{II}$
$O_{III}$
Axial drift
Axial drift
Axial drift
d
$O_{II}$
RIDE-OPM
NFP

**Supplementary Fig. 3 | Comparison of OPM and RIDE-OPM Performance Using Simulated PSFs Under Mechanical Drift During Long-Term Imaging. a,** In conventional OPM, $O_{III}$ must be precisely aligned with the tilted LS plane. When the NFP of $O_{III}$ aligns with the LS, a well-formed PSF is achieved, albeit with some NA loss. **b,** Lateral drift of the $O_{II}$ causes the light to shift off-axis relative to the pupil of $O_{II}$. This misalignment alters the tilt of the image plane, displaces the NFP, and results in both light loss and reduced NA, leading to PSF broadening. **c,** Axial displacement disrupts the sensitive alignment between the LS image and the NFP of $O_{III}$, producing a defocused PSF over time. **d,** RIDE-OPM eliminates the need for $O_{III}$. Even in the presence of minor drift in $O_{II}$, the use of on-axis light reflection preserves proper image formation, maintaining a high-quality PSF with full NA utilization. NPF, nominal focal plane.

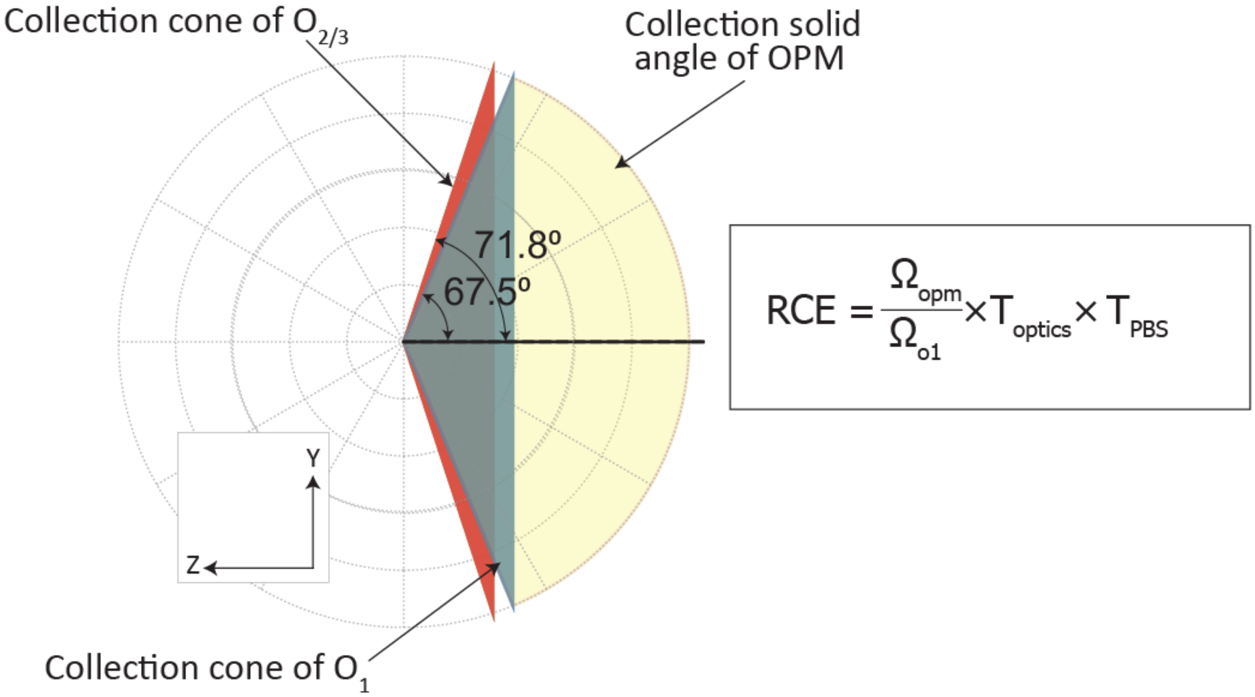


**Supplementary Fig. 4 | Geometric collection efficiency in the RIDE-OPM system.** The geometrical collection efficiency is determined numerically by calculating the area of intersection between the acceptance cones of the objectives on the unit sphere. The red cone represents the collection cone of the secondary objective ($O_2$; NA = 0.95, air), corresponding to a half-angle of ~71.8°, while the blue cone represents the collection cone of the primary objective ($O_1$; NA = 1.4, oil immersion), corresponding to a half-angle of ~67.5°. Both cones are projected into a common coordinate system aligned with the optical axis (black line). The overlapping region, shown in yellow, defines the effective collection solid angle of the system ($\Omega_{OPM}$) and is used to quantify the relative geometric collection efficiency.

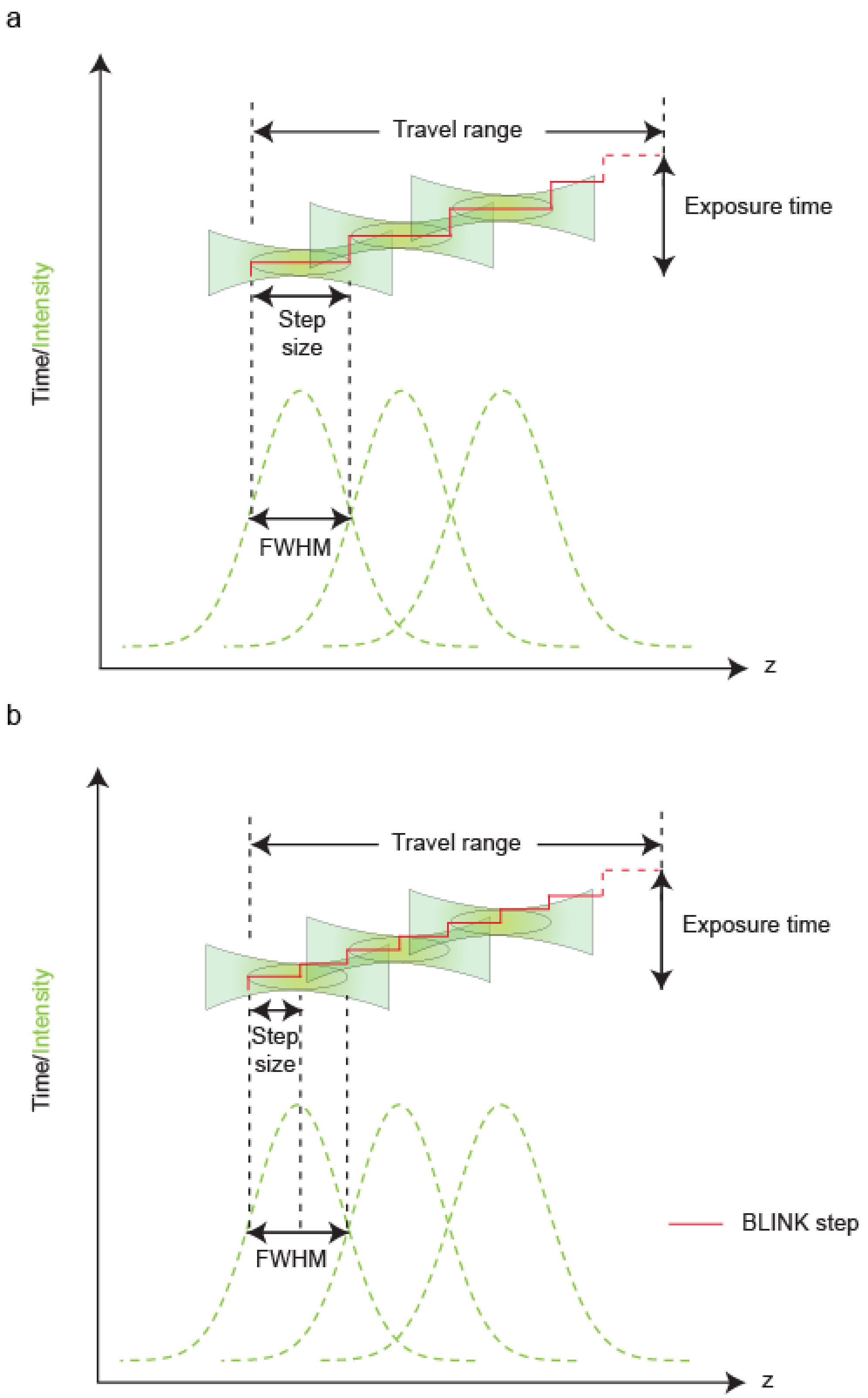


**Supplementary Fig. 5 | Quantification of the step size of the focuser. a-b,** Step size of the focuser during z-depth traversal within the camera's exposure time for capturing an xz' plane. Undersampled step size, which is inadequate for capturing all necessary information (**a**), while Nyquist sampling effectively gathers complete data from the specimen, ensuring detailed imaging (**b**).

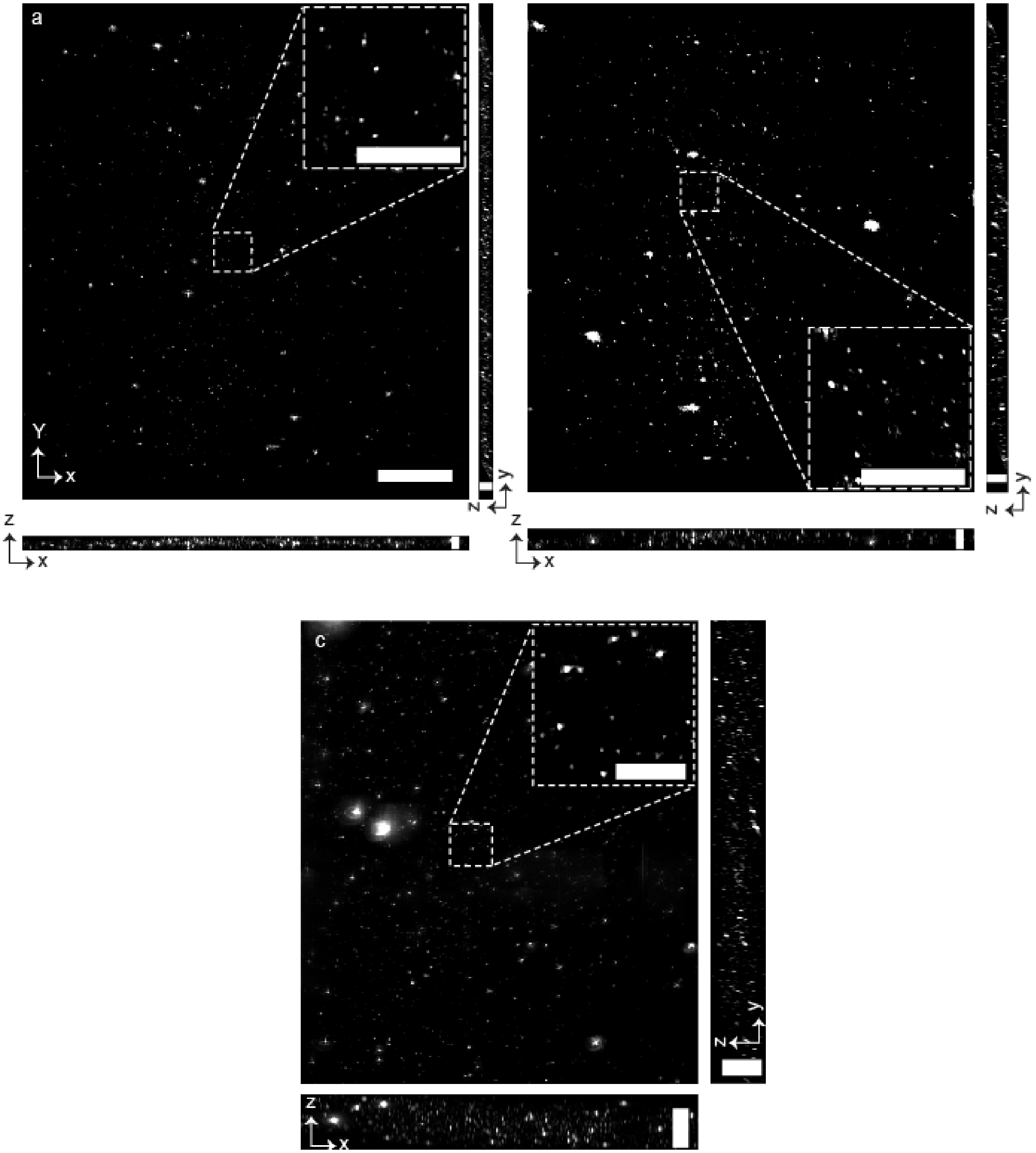


**Supplementary Fig. 6 | Representation of the 3D FOV. a-c,** Maximum intensity projection (MIP) of 100 nm fluorescent beads suspended in glycerol from all three dimensions for scale 1 at 30° of LS tilt (**a**), scale 1 at 45° of LS tilt (**b**) and scale 2 (**c**). The visualization effectively illustrates the three-dimensional field of view (3D FOV) of the beads, highlighting their distribution throughout the sample. Scale bars, 30 μm (xy), 5 μm (xz and yz) (**a**); 30 μm (xy), 8 μm (xz and yz) (**b**); 50 μm (xy), 20 μm (xz and yz) (**c**); 10 μm (inset of **a**, **b**, **c**).

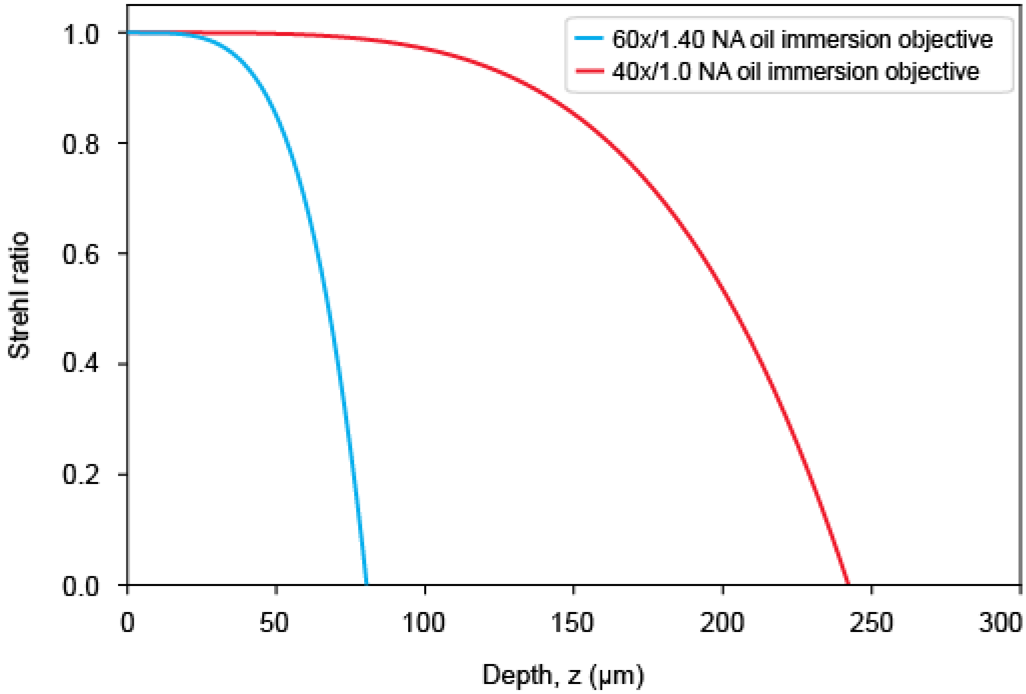


**Supplementary Fig. 7 | Strehl ratio with respect to z-depth.** Strehl ratio plot as a function of z-depth for our primary objective used in scale 1 and scale 2, illustrating the objective's remote focusing performance across various depths[5]. The plot shows that a Strehl ratio of 80% is achievable up to 53 μm for Scale 1 and 162 μm for Scale 2 along the z-axis, confirming that both configurations provide consistent imaging performance within the required 4–30 μm imaging depth range for each respective scale.

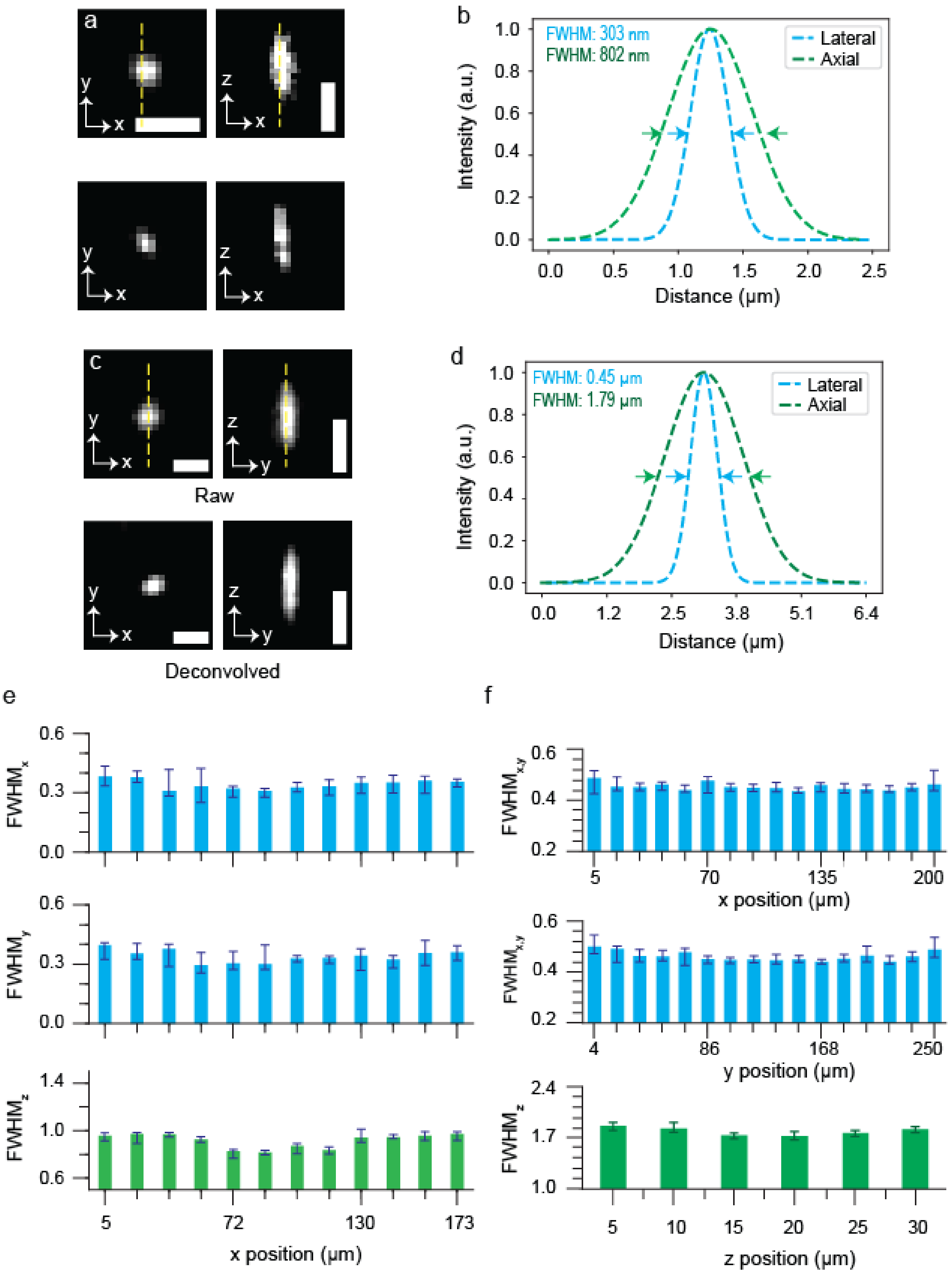
a
y
x
z
x
b
FWHM: 303 nm
FWHM: 802 nm
Lateral
Axial
Intensity (a.u.)
Distance (μm)
1.0
0.8
0.6
0.4
0.2
0.0
0.0
0.5
1.0
1.5
2.0
2.5
c
Raw
Deconvolved
d
FWHM: 0.45 μm
FWHM: 1.79 μm
Lateral
Axial
Intensity (a.u.)
Distance (μm)
0.0
1.2
2.5
3.8
5.1
6.4
e
FWHMx
FWHMy
FWHMz
0.6
0.3
0.0
1.4
1.0
0.6
5
72
130
173
x position (μm)
f
FWHMx,y
FWHMz
0.6
0.4
0.2
2.4
1.7
1.0
5
70
135
200
x position (μm)
4
86
168
250
y position (μm)
5
10
15
20
25
30
z position (μm)

**Supplementary Fig. 8 | Microscope characterization. a-d,** Orthogonal representation of both raw and Richardson-Lucy deconvolved PSF for scale 1 at 45° LS tilt (**a,b**) and scale 2 (**c,d**), obtained by imaging 100 nm of fluorescent beads embedded in glycerol. Resolution was determined from the Gaussian-fitted line profile. **e-f,** Resolution measurements across the entire FOV demonstrated consistent imaging performance throughout for both scale 1 at 45° LS tilt (**e**) and scale 2 (**f**).

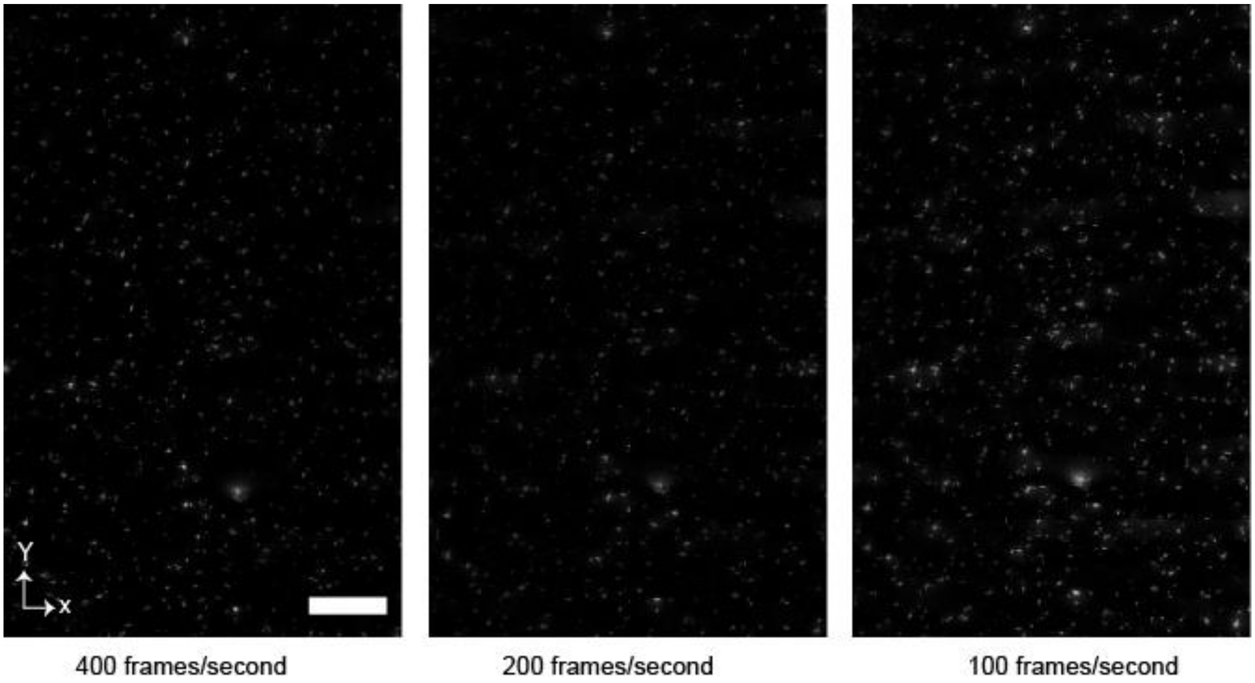


**Supplementary Fig. 9** | **Investigation of imaging speed.** Maximum intensity projection (MIP) of 500 nm fluorescent beads embedded in glycerol, imaged at various frame rates. The sequence demonstrates the microscope's ability to achieve camera-limited imaging speeds even for 30 μm of depth above the cover slip, showcasing how different speeds impact the clarity and quality of the images. Scale bar, 30 μm.

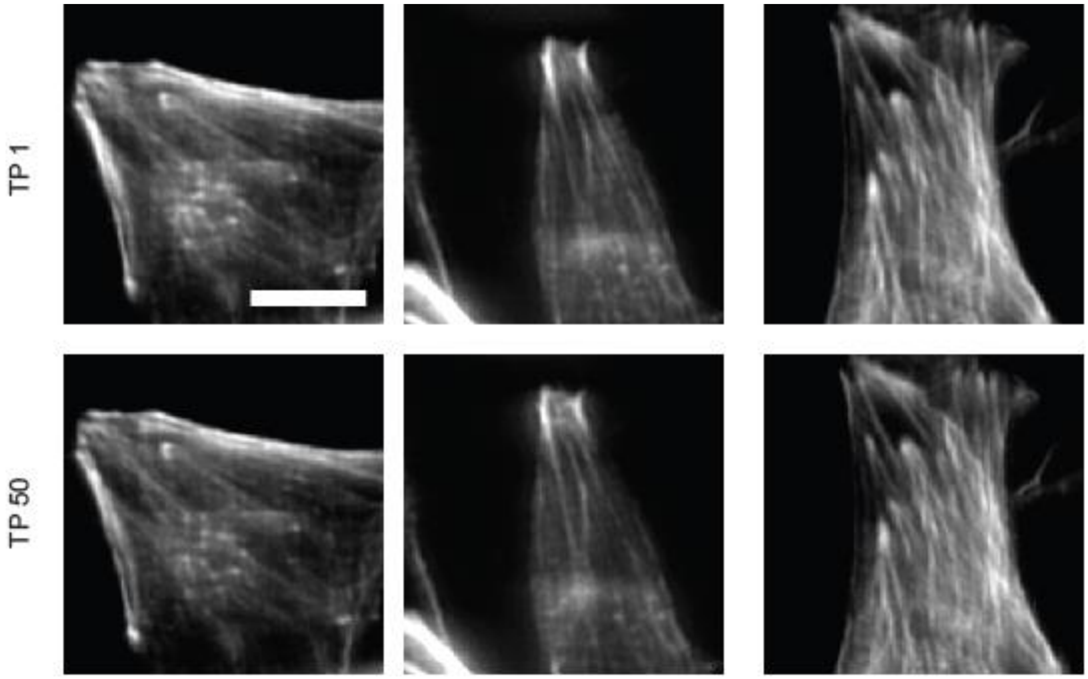


**Supplementary Fig. 10 | Hour-long cellular imaging using ASLM for photobleaching analysis.** MDA-MB-231 human triple-negative breast cancer cells, stained for F-actin with phalloidin–Alexa Fluor 647, were imaged over a one-hour period by ASLM microscope. Fluorophore bleaching was quantified across multiple regions of interest (ROIs) over 50 time points acquired during the experiment, and a comparative analysis of bleaching across different imaging modalities is presented in Fig. 2h. Scale bar, 10 μm.

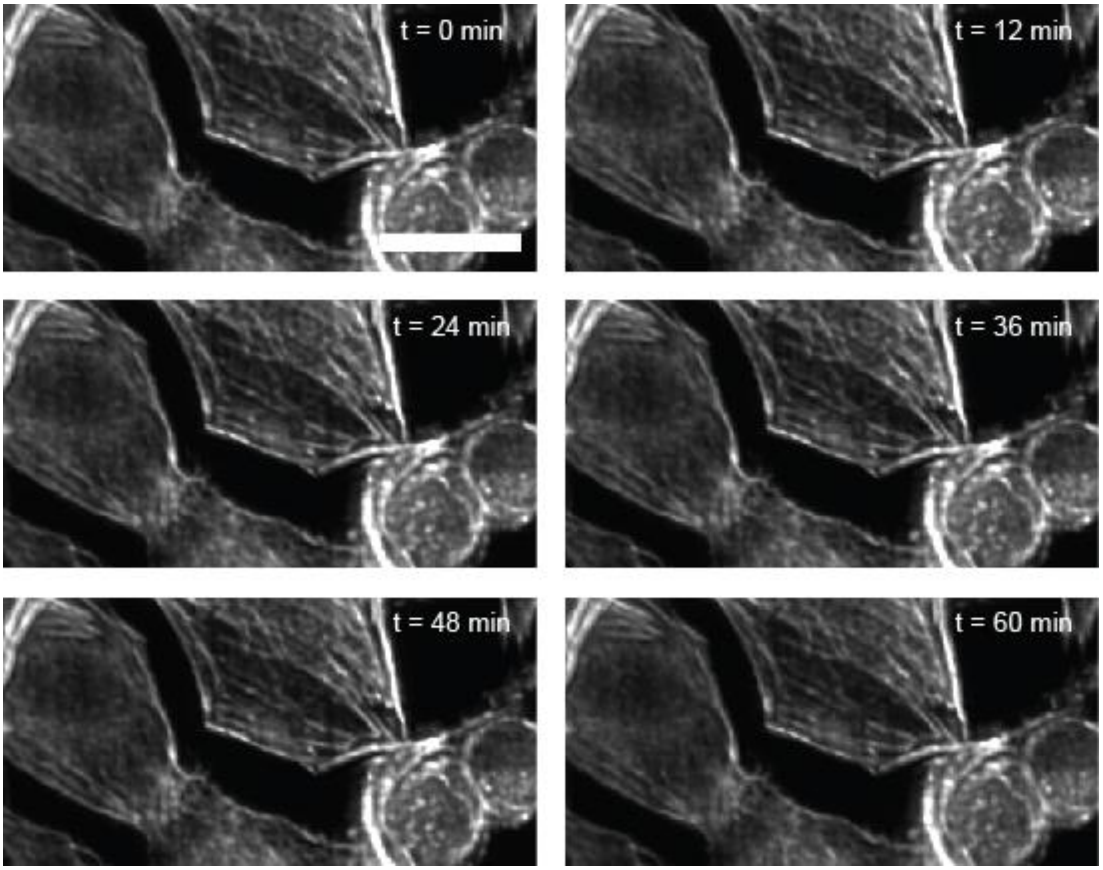


**Supplementary Fig. 11 | Hour-long cellular imaging using RIDE-OPM for photobleaching analysis.** MDA-MB-231 human triple-negative breast cancer cells, stained for F-actin with phalloidin–Alexa Fluor 647, were imaged over a one-hour period by RIDE-OPM system. Fluorophore bleaching was quantified across multiple regions of interest (ROIs) over 50 time points acquired during the experiment, and a comparative analysis of bleaching across different imaging modalities is presented in Fig. 2h. Scale bar, 20 μm.

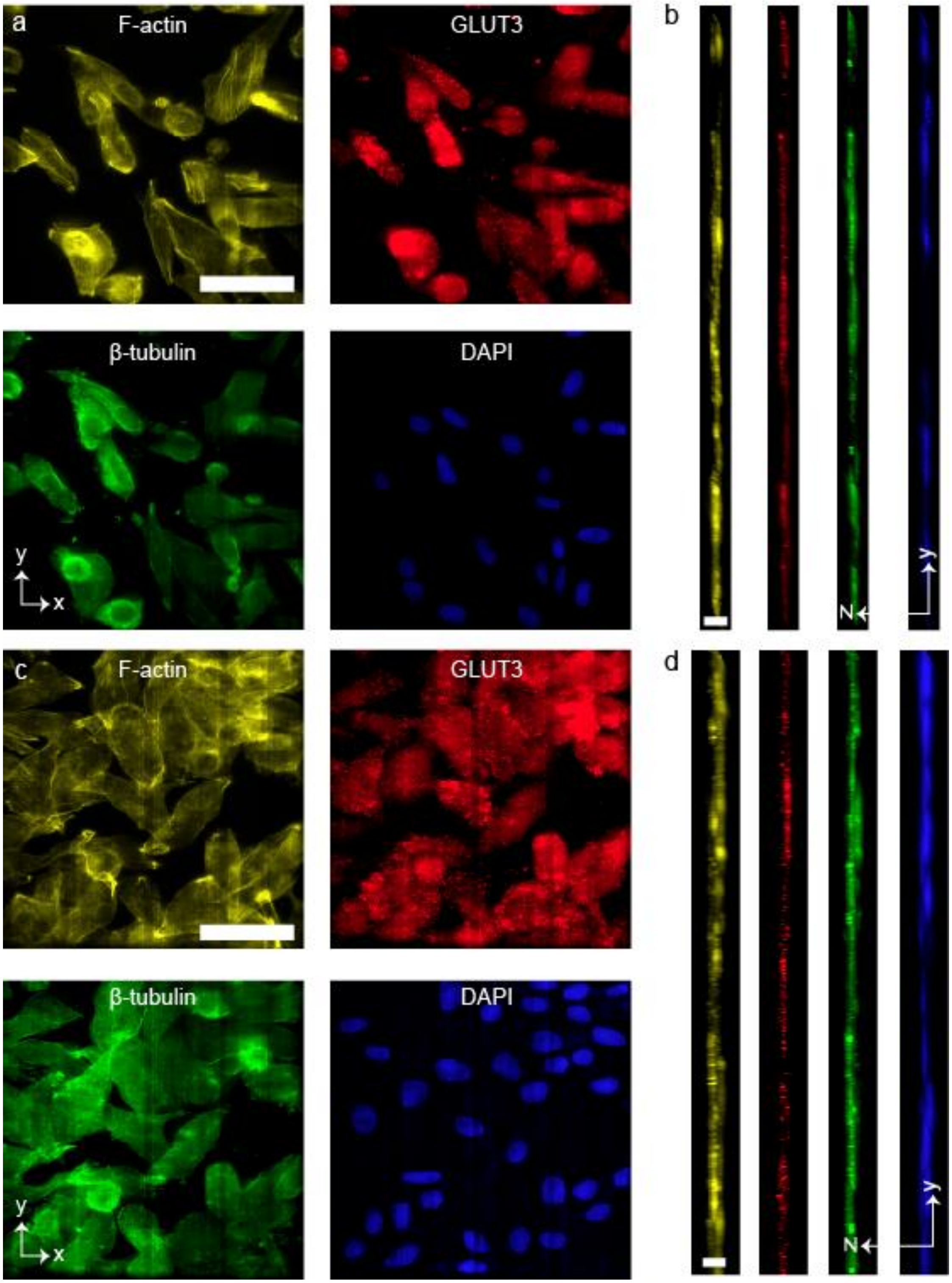


**Supplementary Fig. 12 | Tilt-independent multi-color imaging of human breast cancer cell. a-d,** Orthogonal maximum intensity projection (MIP) of MDA-MB-231 (human triple-negative breast cancer) cell, imaged with 30° (**a-b**) and 45° (**c-d**) LS tilt angles. The cell was stained for F-actin (yellow), GLUT3 (red), β-tubulin (green), and DAPI (blue; nuclei). Scale bar, 50 µm (**a,c**), 6 µm (**b,d**).

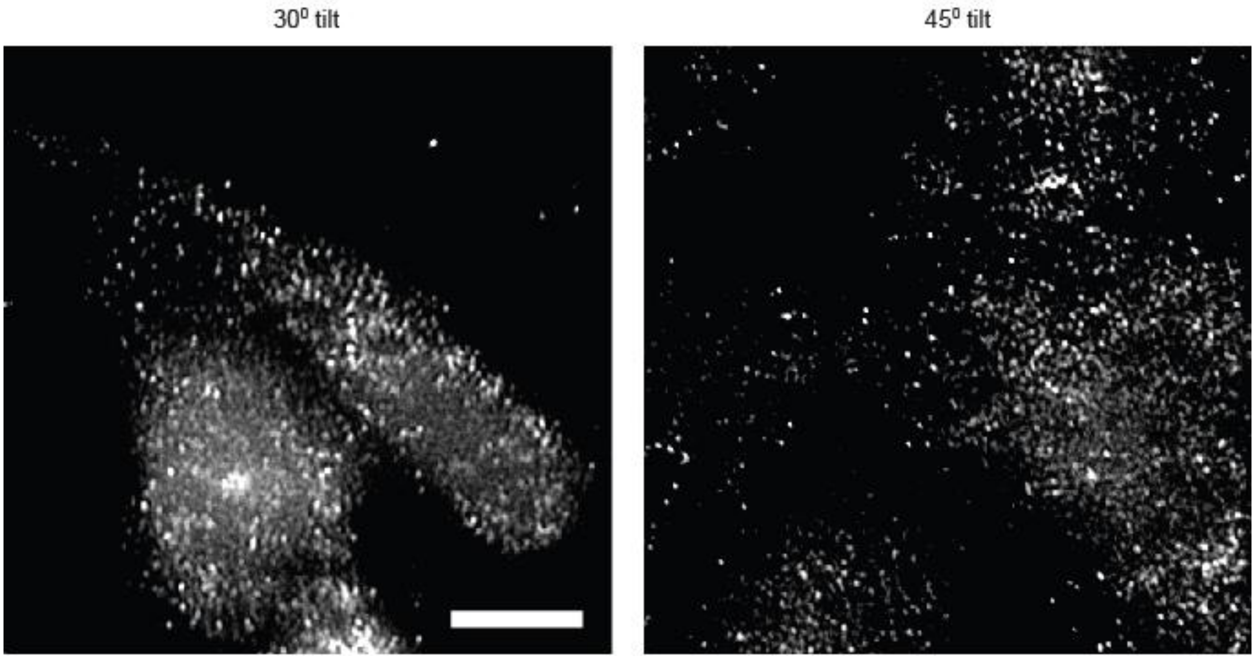


**Supplementary Fig. 13 | Tilt-independent imaging.** GLUT3 distribution in malignant breast epithelial cells (MDA-MB-231, human triple-negative breast cancer) was imaged at light-sheet tilt angles of 30° and 45°. GLUT3 labeling was performed using a primary anti-GLUT3 antibody (1:200), followed by an Alexa Fluor 561–conjugated secondary antibody. Scale bars, 10 μm.

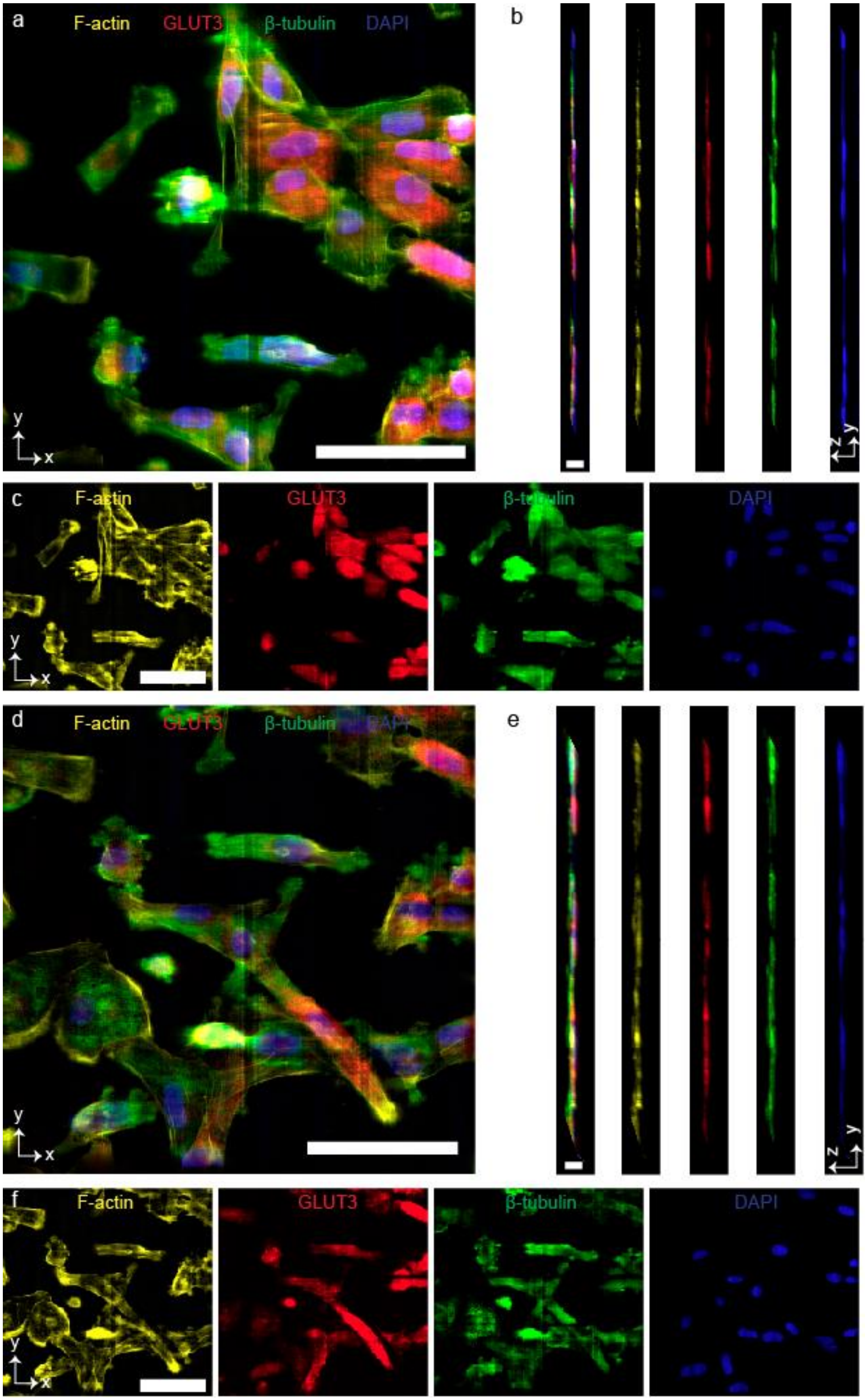
a
F-actin
GLUT3
β-tubulin
DAPI
y
x
b
z
y
c
F-actin
GLUT3
β-tubulin
DAPI
d
F-actin
GLUT3
β-tubulin
DAPI
e
f
F-actin
GLUT3
β-tubulin
DAPI

**Supplementary Fig. 14 | Tilt-independent multi-color imaging of human mammary epithelial cell. a-f,** Orthogonal maximum intensity projection (MIP) of MCF10A (human non-tumorigenic mammary epithelial) cell, imaged with 30° (**a-c**) and 45° (**d-f**) LS tilt angles. The cell was stained for F-actin (yellow), GLUT3 (red), β-tubulin (green), and DAPI (blue; nuclei). Scale bar, 50 μm (**a,c**), 6 μm (**b,d**).

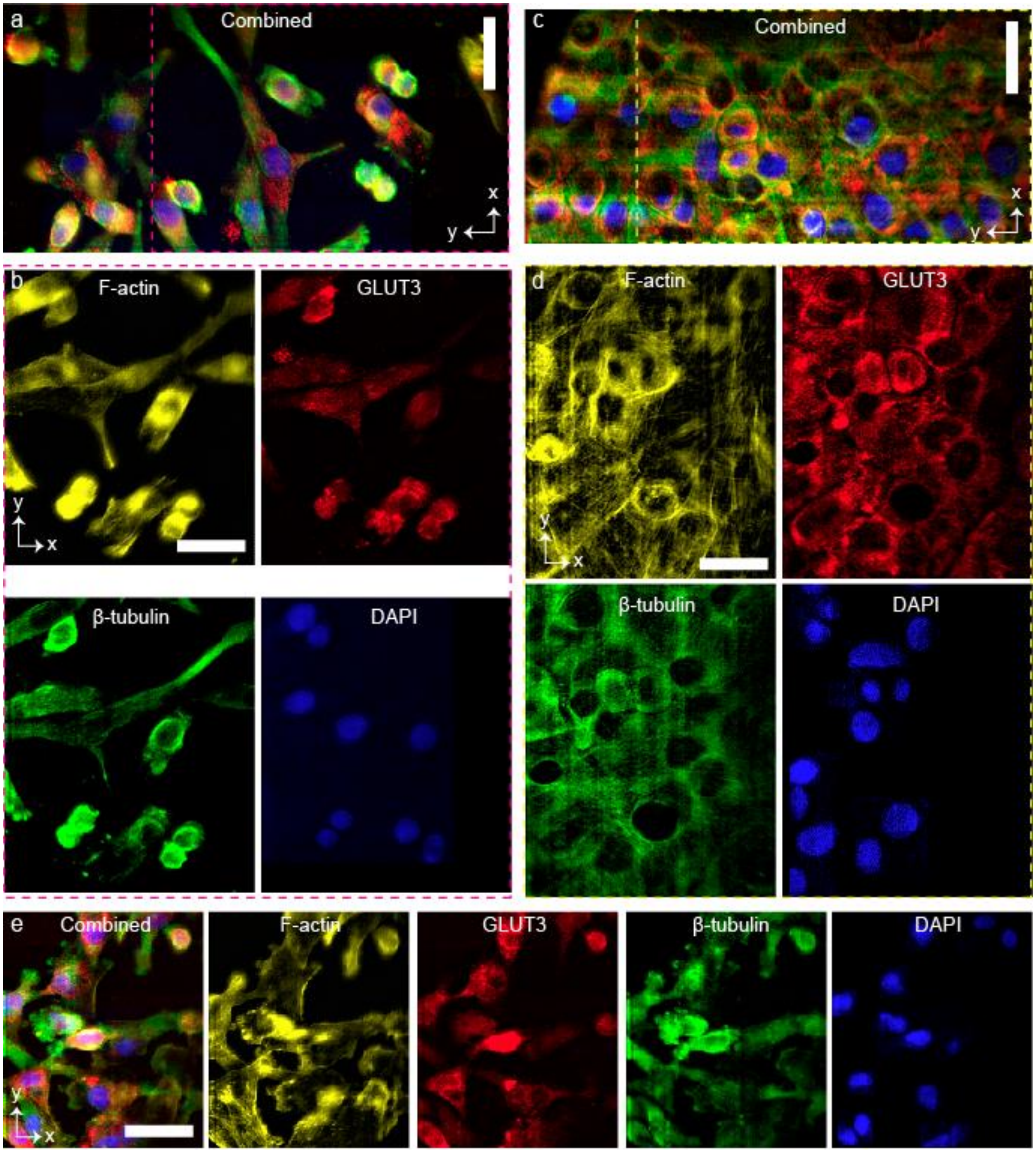


**Supplementary Fig. 15 | Multi-color imaging of GLUT3, F-actin, and β-tubulin in normal and malignant breast epithelial cell lines. a-e,** MDA-MB-231 (human triple-negative breast cancer) cell (**a-b)**, MCF10A (human non-tumorigenic mammary epithelial) cell (**c-d**), EO771 (mouse triple-negative breast cancer) cell (**e**) stained for F-actin (yellow), GLUT3 (red), β-tubulin (green), and DAPI (blue; nuclei). F-actin was visualized using phalloidin-Alexa Fluor 647. GLUT3 staining was performed using a primary antibody (anti-GLUT3, 1:200) and Alexa Fluor 561-conjugated secondary antibody. β-tubulin was labelled with an anti-β-tubulin antibody (1:500) and Alexa Fluor 488 secondary antibody. Scale bars, 30 μm (**a-e**).

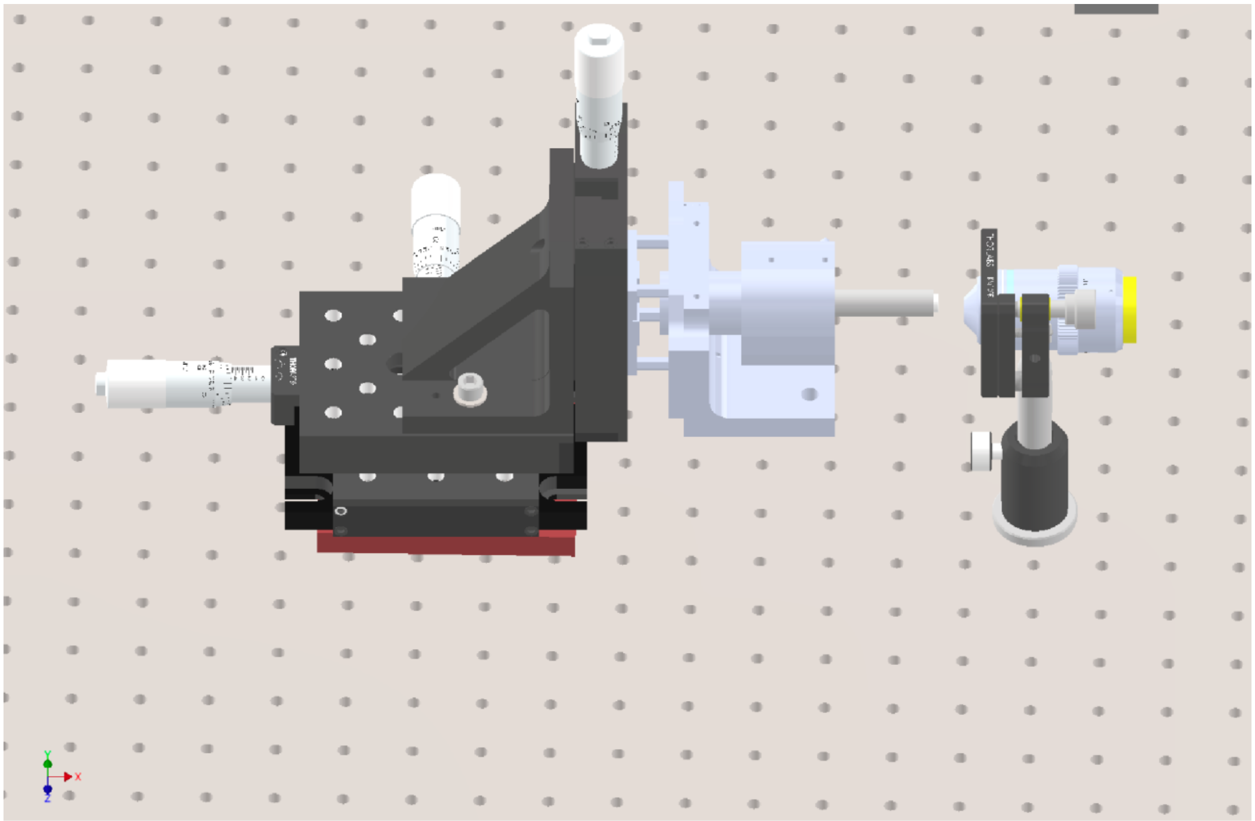

**Supplementary Fig. 16 | Secondary objective and the focuser.** 3D layout of the secondary objective and the focuser (LFA, Equipment solutions/BLINK, Thorlabs) with a 7 mm diameter mirror attached to its shaft.

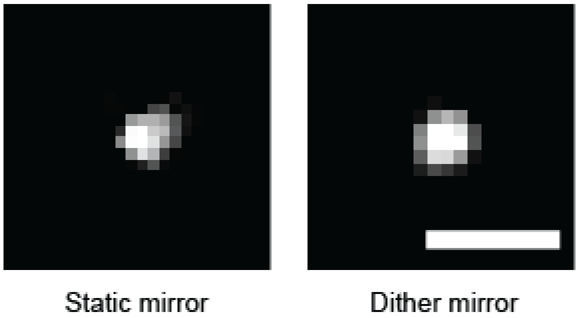


**Supplementary Fig. 17 | PSF comparison between the platforms based on static and dithered mirror.** Comparison of the PSFs obtained with static and dithered mirror configurations shows similar profile characteristics, indicating equivalent performance in terms of spatial resolution. Scale bar, 1 μm.

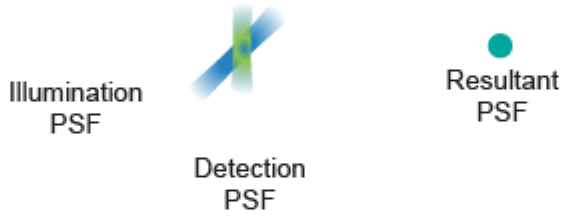


**Supplementary Fig. 18 | RIDE-OPM system PSF.** Due to the optical sectioning offered by the LS illumination, the overall system PSF (right) is the product of the illumination PSF (blue) and the detection PSF (green) shown on the left.

a

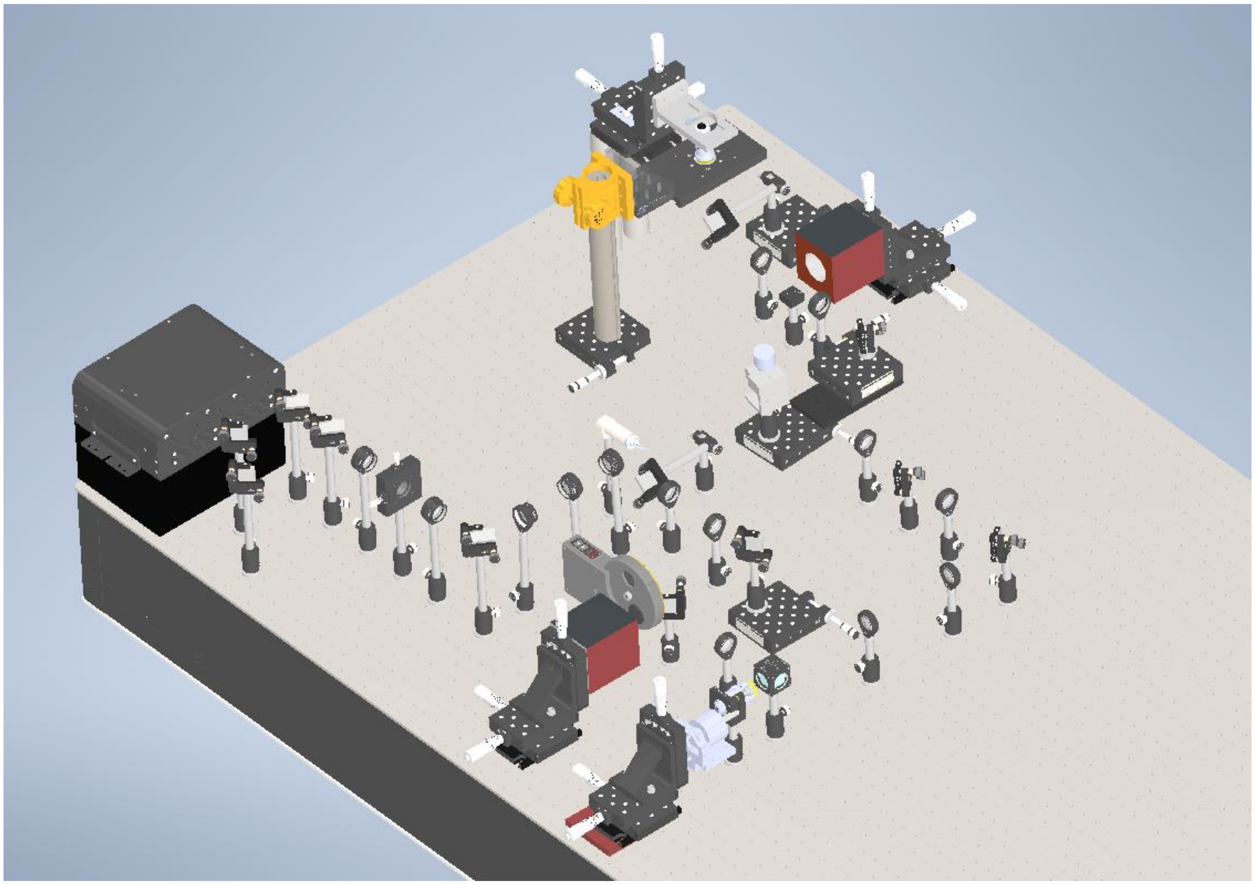

b

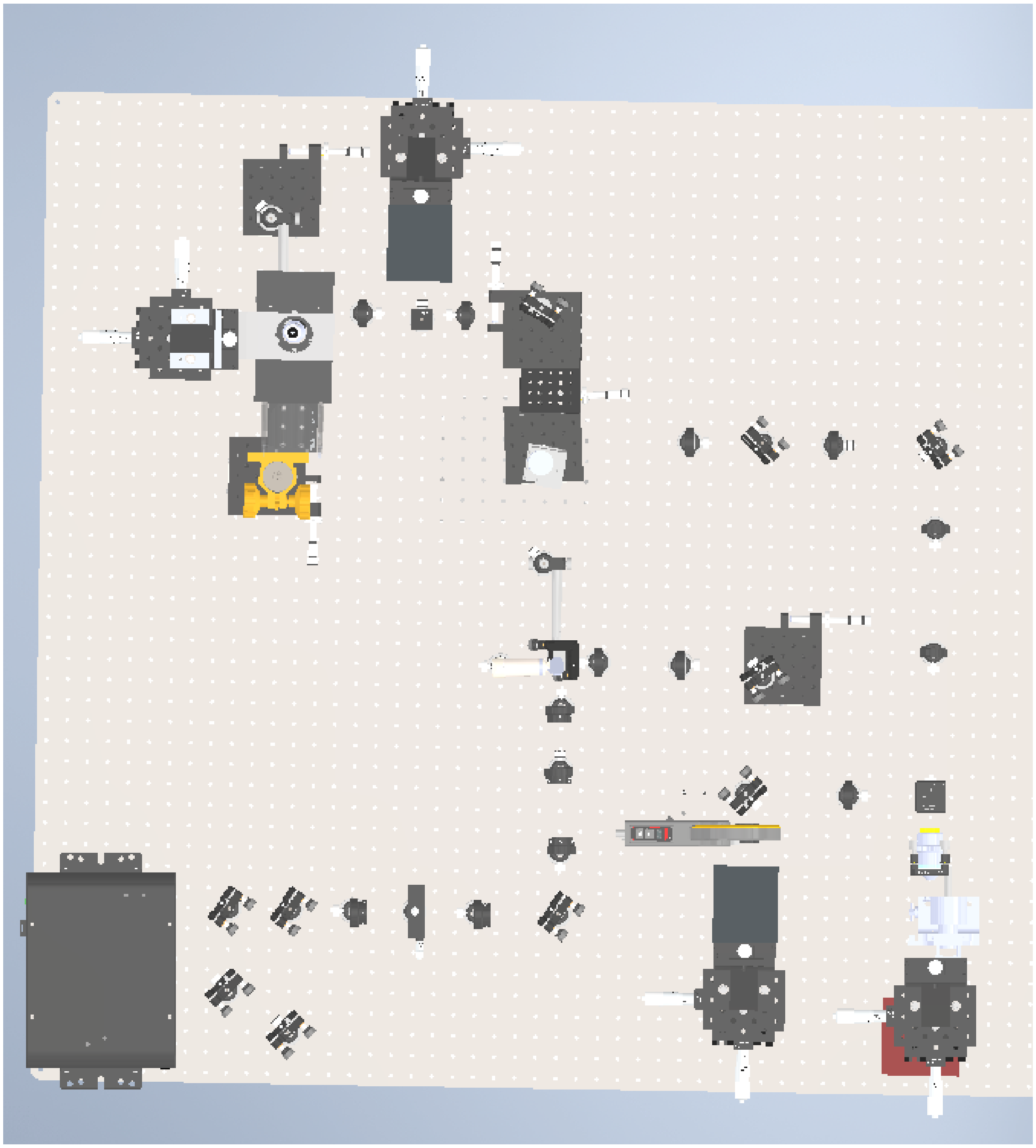

**Supplementary Fig. 19i** | **Microscope layout.** 3D visualization of the RIDE-OPM microscope (scale 1) layout at an angle (**a**) and from top (**b**). The essential equipment required for this setup is detailed in **Supplementary Table 2a**. This visualization helps in understanding the spatial arrangement and integration of the various components that make up the RIDE-OPM system.

a

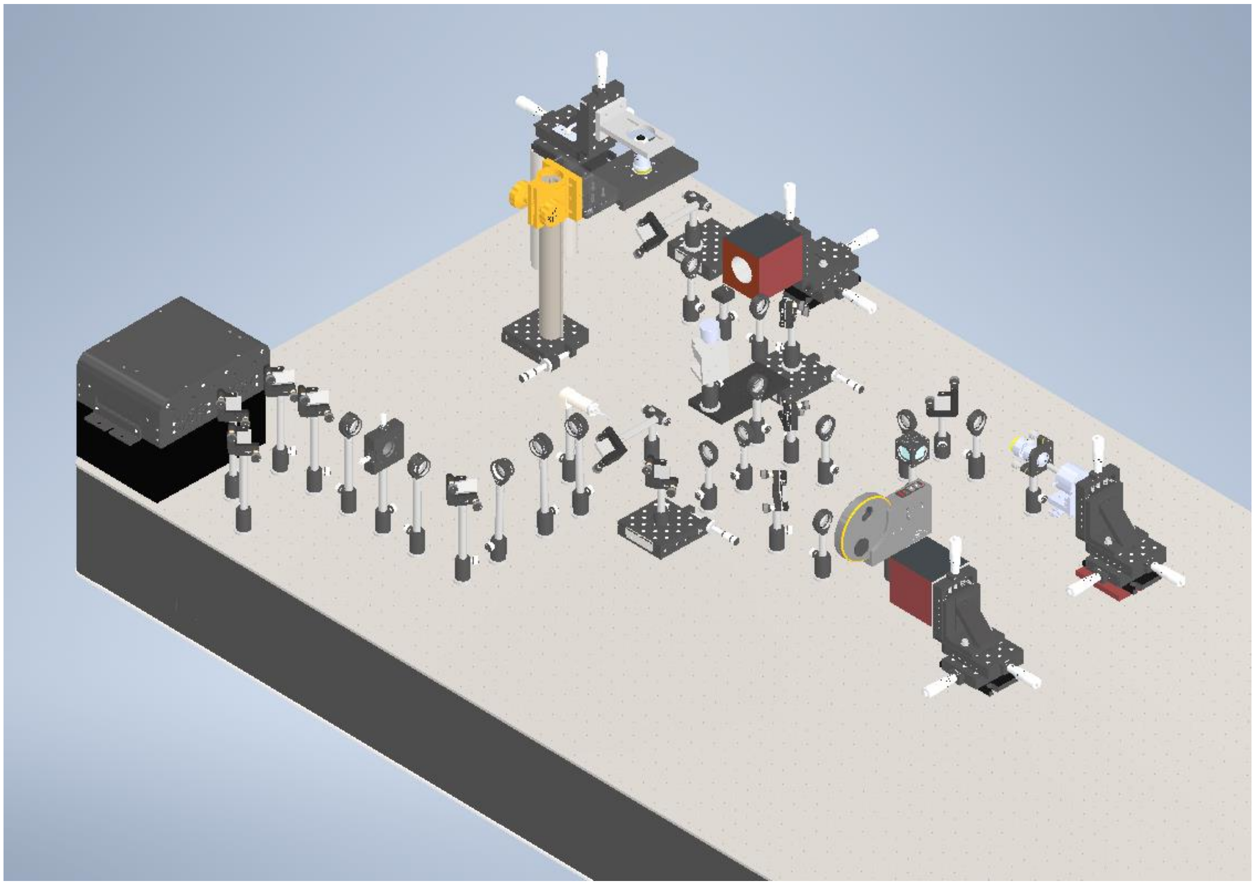

b

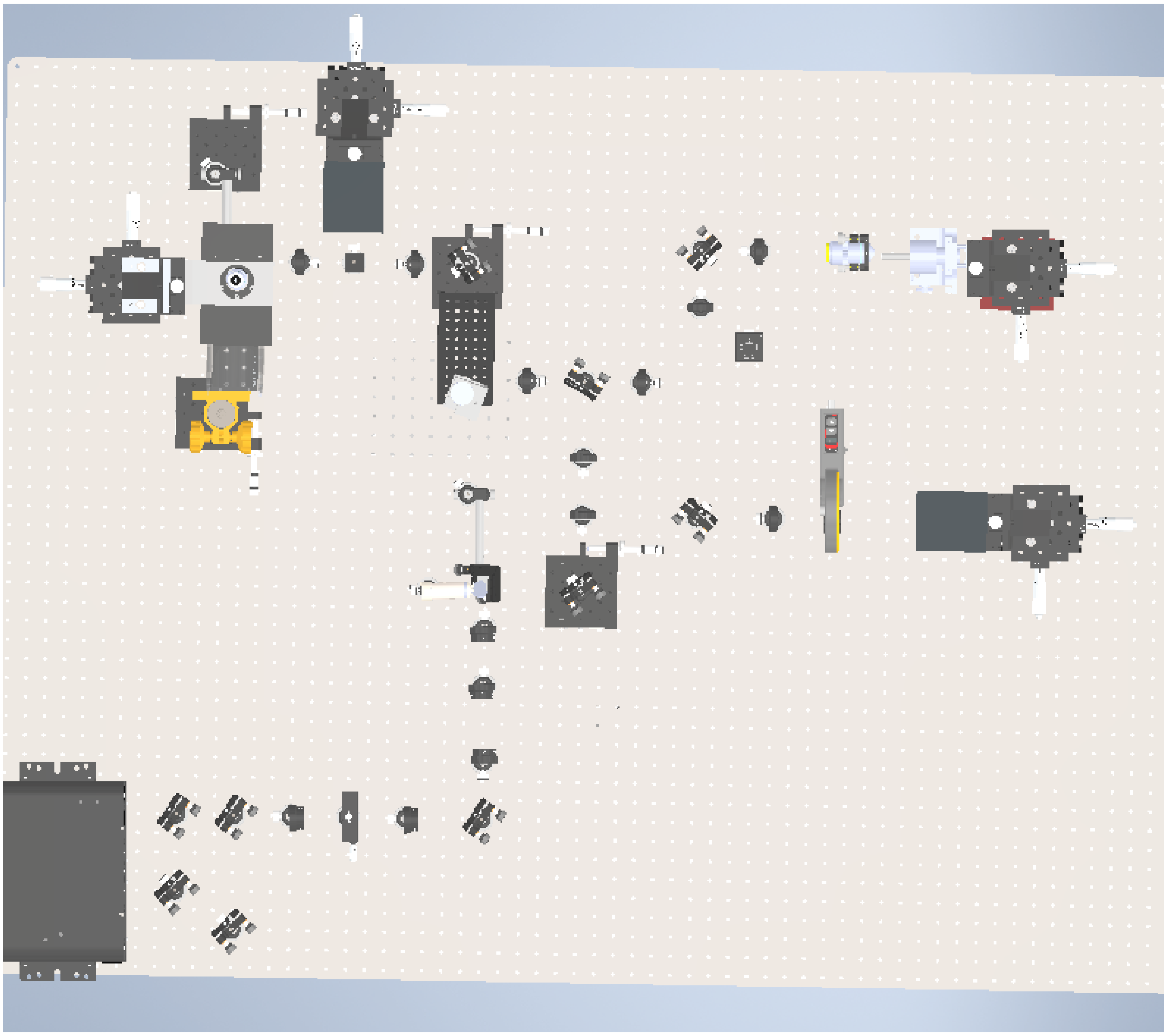

**Supplementary Fig. 19ii** | **Microscope layout.** 3D visualization of the RIDE-OPM microscope (scale 2) layout at an angle (**a**) and from top (**b**). The essential equipment required for this setup is detailed in **Supplementary Table 2b**. This visualization helps in understanding the spatial arrangement and integration of the various components that make up the RIDE-OPM system.

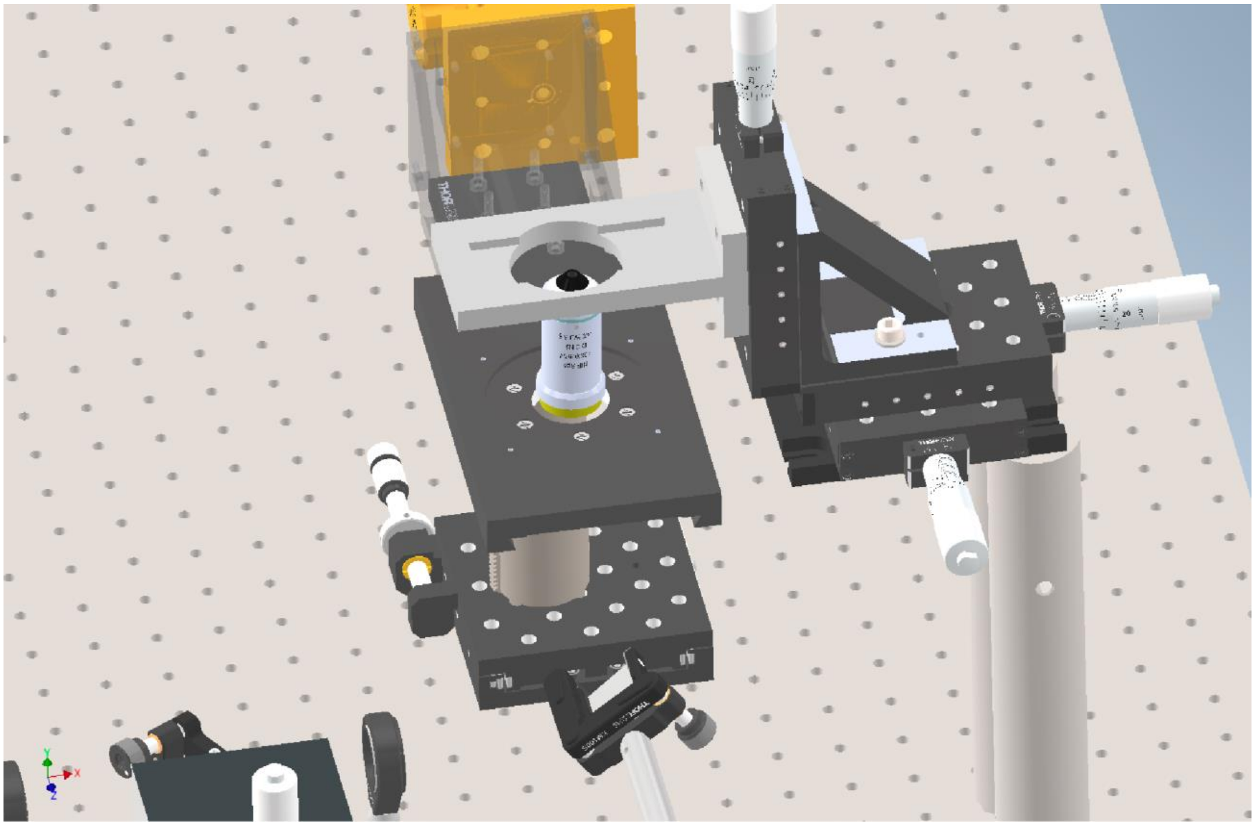

**Supplementary Fig. 20** | **Primary Objective and open-top sample mounting.** 3D layout of primary objective with a simple sample mounting stage for open-top configuration (3D motorized stage for sample translation is not shown here).

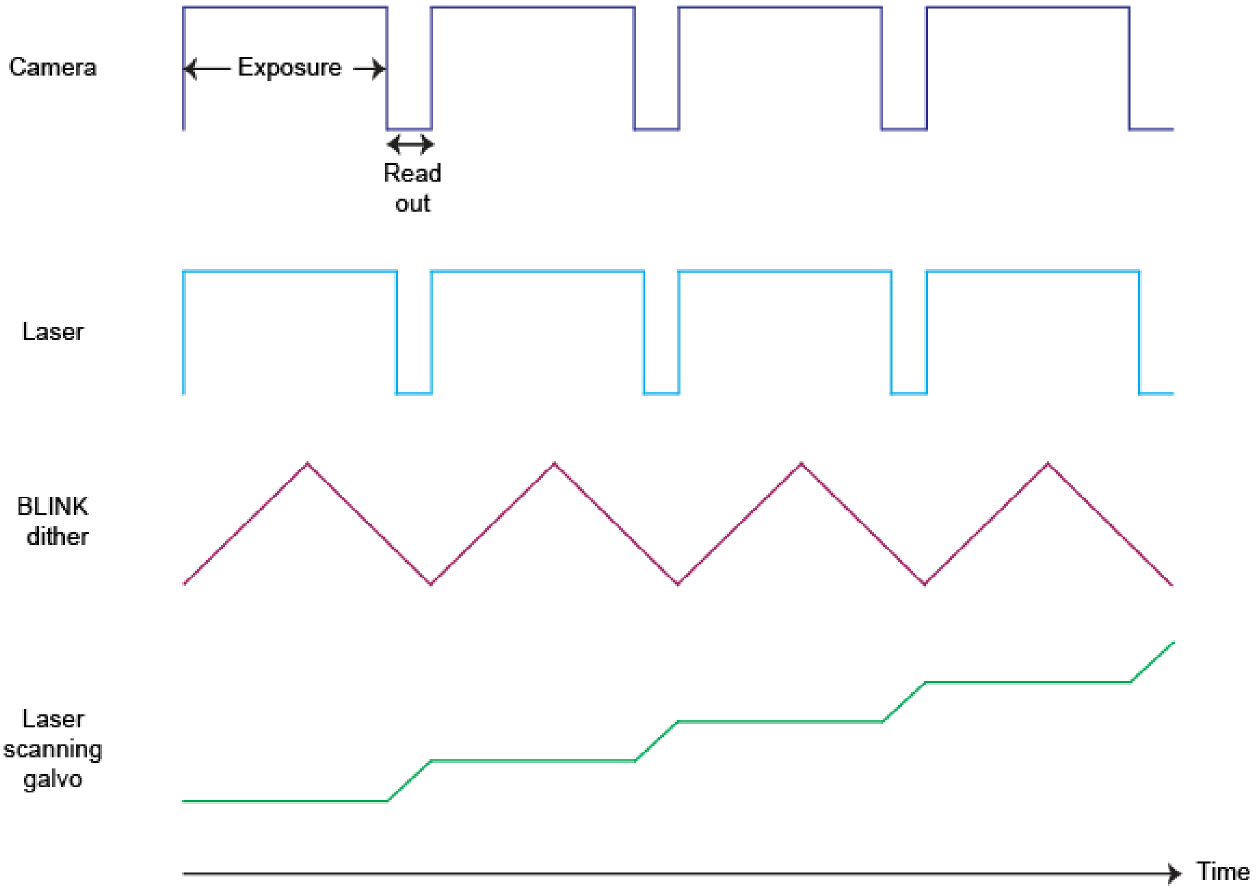


**Supplementary Fig. 21 | Timing diagram.** The timing diagram displays the synchronization of operating voltages and input signals for different components of the microscope in relation to the camera frame acquisition time, for a single channel imaging modality. The diagram outlines the precise timing sequences necessary to ensure coordinated operation and optimal image capture, highlighting the interplay between various system elements and the timing of each relative to the camera's frame capture.

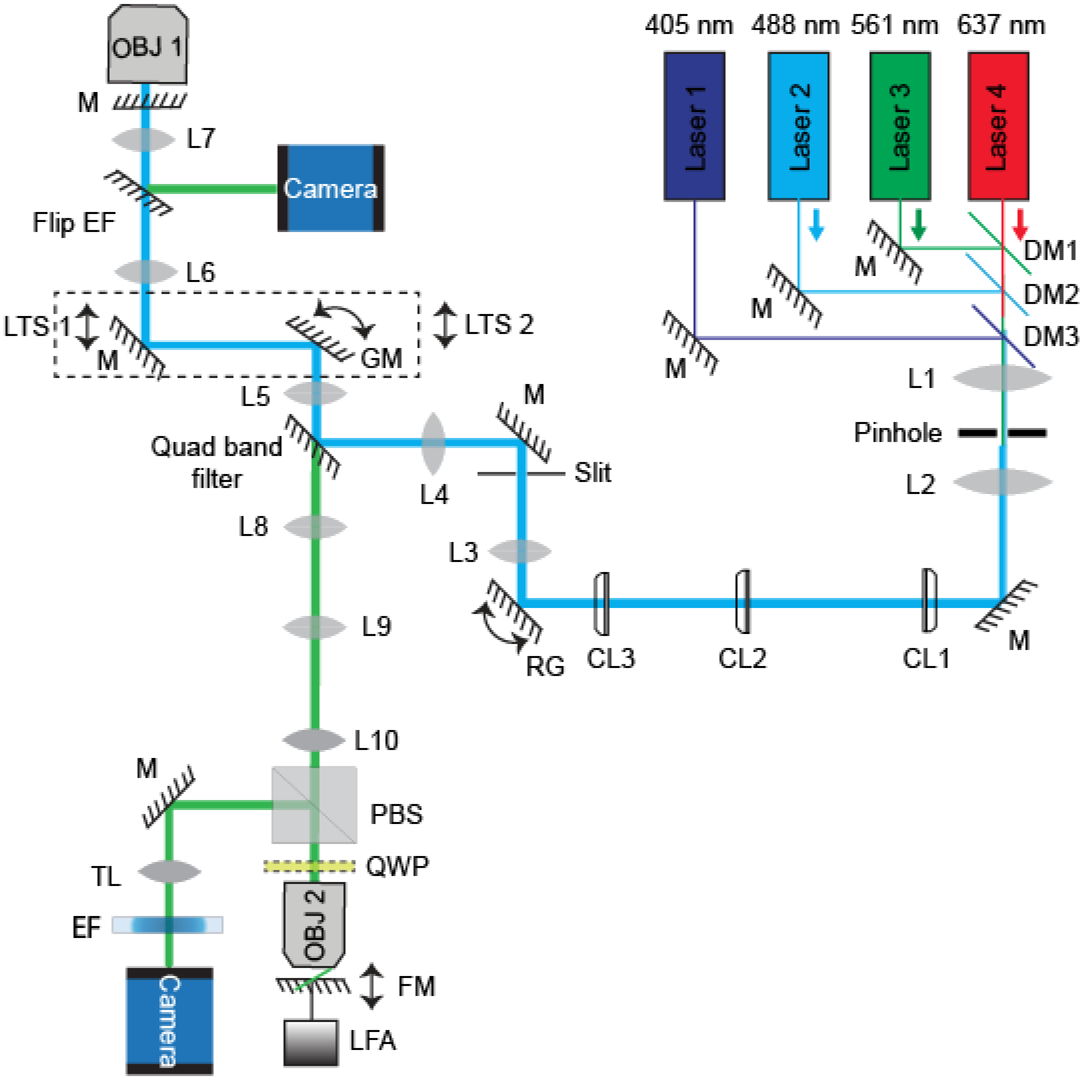


**Supplementary Fig. 22i** | **Schematic diagram of RIDE-OPM scale 1.** A comprehensive view of the RIDE-OPM system's optical setup for scale 1. It includes all the necessary components, which are detailed in **Supplementary Table 2a**, offering a clear understanding of the system configuration and component interrelationships within the proposed RIDE-OPM system.

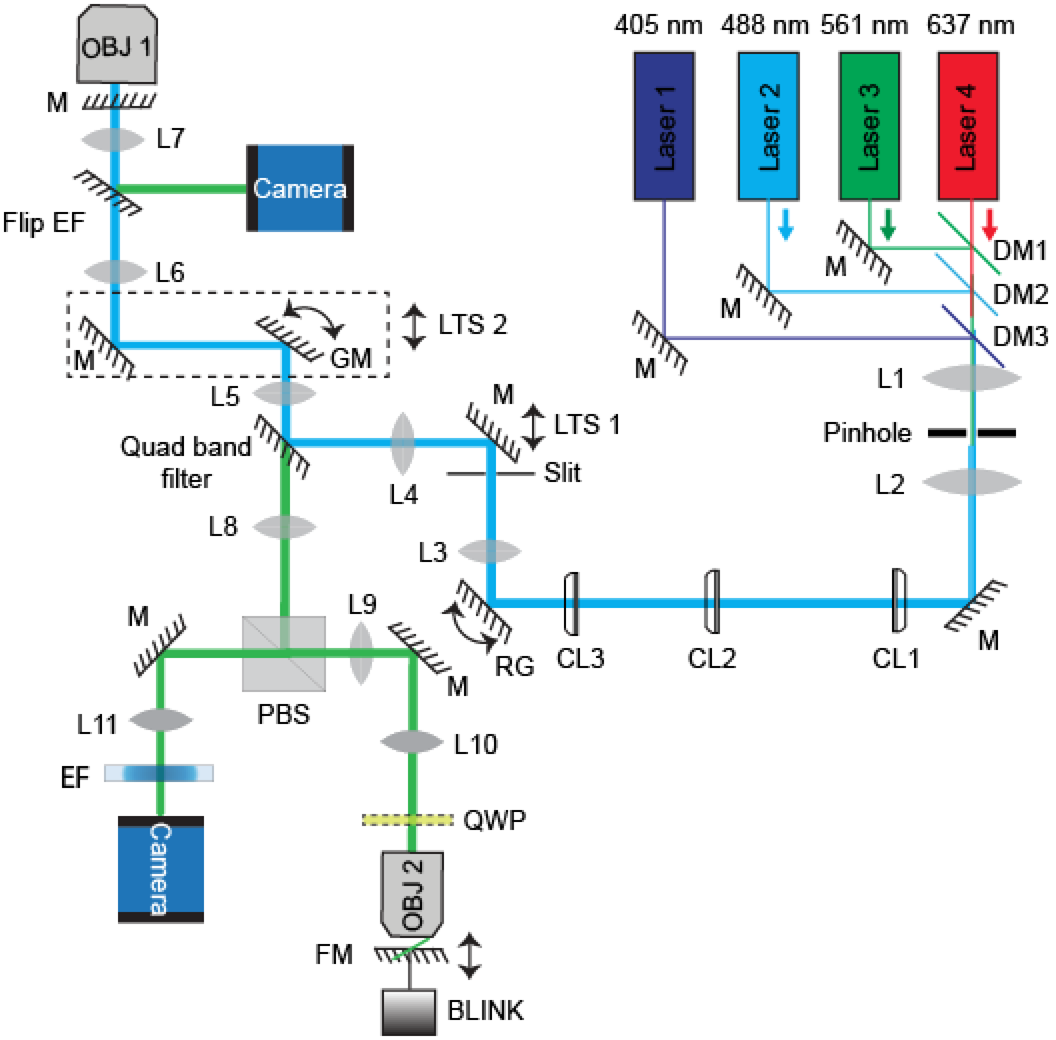


**Supplementary Fig. 22ii** | **Schematic diagram of RIDE-OPM scale 2.** A comprehensive view of the RIDE-OPM system's optical setup for scale 2. It includes all the necessary components, which are detailed in **Supplementary Table 2b**, offering a clear understanding of the system configuration and component interrelationships within the proposed RIDE-OPM system.

### Supplementary Tables

**Supplementary Table 1 |** Detailed comparison of methods aiming to capture full NA.

| Items | RIDE-OPM | obSTORM[6] | dOPM[7,8] | Full-aperture OPM[9] |
|---|---|---|---|---|
| Primary objective | Nikon (60×), Oil immersion (Scale 1)<br>Nikon (40×), Oil immersion (Scale 2) | UPLSAPO 60XW Olumpus | Nikon (40×), water immersion[a]<br>Nikon (60×), water immersion[b] | Olympus (60×), water-dipping<br>Olympus (20×), water-dipping |
| Primary objective NA | 1.40 (Scale 1)<br>1.0 (Scale 2) | 1.20 | 1.15[a]<br>1.20[b] | 1.10<br>0.50 |
| Scanning-descanning modality | Yes | No | Yes | Yes |
| Volumetric imaging | Yes | No | Yes | Yes |
| NA utilization | 100% | Loses NA | | 100% |
| Lateral Field of view (FOV) | 173 µm × 200 µm (Scale 1)<br>200 µm × 252 µm (Scale 2) | | 256 µm × 256 µm[a]<br>100 µm × 100 µm[b] | 240 µm × 420 µm (60×)<br>720 µm × 540 µm (20×) |
| Lateral resolution | ~ 300 nm (Scale 1)<br>~ 440 nm (Scale 2) | ~ 400 nm<br>~ 44 (with SMLM*) | ~ 480 nm[a]<br>~ 300 nm[b] | ~ 600 nm (NA 1.1)<br>~ 1.71 µm (NA 0.5) |
| Stability | Stable<br>(May appear little vulnerability to voice coil) | Stable | Highly vulnerable to the stage movement | Highly vulnerable due to the ETL** |
| Imaging speed | Camera limited | Camera limited | 90 fps*** | |
| Computational requirement | Deskew | No | Fusion + Deskew | Deskew |

* Single-molecule super resolution microscopy
** Electro-tunable lens
*** Frames per second
[a] H. Sparks et al.[7]
[b] H. Sparks et al.[8]

**Supplementary Table 2a | Equipment list.** Detail list of materials used to construct RIDE-OPM scale 1.

| Sl. No. | Description | Item abbreviation | Part number | Part number company | Qty | Remarks |
|---|---|---|---|---|---|---|
| 1 | 4 channel combo OBIS laser | Laser 0 | LX 637-140C | Coherent | 1 | |
| | | Laser 1 | LX 488-50C | | | |
| | | Laser 2 | LX 561-50 | | | |
| | | Laser 3 | LX 405 | | | |
| 2 | 613 nm Dichroic beam splitter | DM1 | LM01-613-25 | Semrock | 1 | |
| 3 | 503 nm Dichroic beam splitter | DM2 | LM01-503-25 | Semrock | 1 | |
| 4 | 445 nm Dichroic beam splitter | DM1 | DMLP445 | Thorlabs | 1 | |
| 5 | f = 50 mm, Ø1" achromatic doublet | L1, L2 | AC254-50-A-ML | Thorlabs | 2 | |
| 6 | f = 150 mm, Ø1" achromatic doublet | L3, L4 | AC254-150-A-ML | Thorlabs | 2 | |
| 7 | f = 75 mm, Ø1" achromatic doublet | L5, L8 | AC254-75-A-ML | Thorlabs | 2 | |
| 8 | f = 100 mm, Ø1" achromatic doublet | L6, L9 | AC254-100-A-ML | Thorlabs | 2 | |
| 9 | f = 200 mm, Ø2" achromatic doublet | L7, L10 | ACT508-200-A-ML | Thorlabs | 2 | |
| 10 | f = 200 mm, Tube lens | TL | TTL-200 | Thorlabs | 1 | |
| 11 | 50 µm pinhole | Pinhole | P50D | Thorlabs | 1 | |
| 12 | Polarizing beam splitter | PBS1 | 10FC16PB.7 | Newport Corporation | 2 | |
| 13 | f = 25 mm, Ø0.5" cylindrical achromat | CL1 | 68-160 | Edmund Optics | 1 | |
| 14 | f = 200 mm, Ø1" cylindrical achromat | CL2 | ACY254-200-A | Thorlabs | 1 | |
| 15 | f = 50 mm, Ø1" cylindrical achromat | CL3 | ACY254-50-A | Thorlabs | 1 | |
| 16 | Resonant mirror galvanometer | Resonant mirror | CRS 4 kHz | Cambridge Technology | 1 | |
| 17 | Adjustable mechanical slit | Slit | VA100 | Thorlabs | 1 | |
| 18 | Galvanometric mirror | Galvo | GVS011 | Thorlabs | 1 | |
| 19 | Quarter wave plate | QWP | AQWP3 | Bolder Vision Optik | 1 | |
| 20 | High speed focuser (Linear focus actuator) | FM | LFA-2010 | Equipment Solutions | 1 | |
| 21 | Microscope objective (60x/NA 1.4 oil) | OBJ1 | Oil immersion objective | Nikon | 1 | |
| 22 | Microscope objective (40x/NA 0.95 air) | OBJ2 | Dry objective | Nikon | 1 | |
| 23 | 12V DC power supply | 12V DC | A12MT400 | Acopian | 1 | |
| 24 | Quad band filter | Quad band filter | ZT 405/488/561 /640rpcv2-UF1 | Chroma | 1 | |
| 25 | Emission filter | Flip EF | FF01-525/30-25 | Semrock | 1 | |
| 26 | Emission filter | EF1 | BLP01-405R-25 | Semrock | 1 | |
| 27 | Emission filter | EF2 | FF01-525/45-25 | Semrock | 1 | |
| 28 | Emission filter | EF3 | FF01-593/40-25 | Semrock | 1 | |

| Sl. No. | Description | Item abbreviation | Part number | Part number company | Qty | Remarks |
|---|---|---|---|---|---|---|
| 29 | Emission filter | EF4 | BLP01-647R-25 | Semrock | 1 | |
| 30 | 1" x 1" protected silver mirror | M | PFSQ10-03-P01 | Thorlabs | few | |
| 31 | 2" x 2" protected silver mirror | M | PFSQ20-03-P01 | Thorlabs | few | |
| 32 | Ø1" protected silver mirror | M | PF10-03-P01 | Thorlabs | few | |
| 33 | Ø7.0 mm protected silver mirror | FM | PF03-03-P01 | Thorlabs | 1 | |
| 34 | single axis translation stage | Lin Tran stage | LT1 | Thorlabs | few | |
| 35 | 6 position motorized filter wheel | Wheel | FW102C | Thorlabs | 1 | |
| 36 | Digital sCMOS camera | Camera | Orca Flash 4.0, V2 model: C13440-20CU | Hamamatsu Corporation | 1 | |
| 37 | Self-Contained XYZ translation stage | 3D Tran stage | LX30 | Thorlabs | 2 | |
| 38 | 3D motorized stage | Mot stage | Model: MP-285A, PCIe 80 7852R | National Instruments | 1 | |

Qty. quantity

**Supplementary Table 2b | Equipment list.** Detail list of materials used to construct RIDE-OPM scale 2.

| Sl. No. | Description | Item abbreviation | Part number | Part number company | Qty | Remarks |
|---|---|---|---|---|---|---|
| 1 | 4 channel combo OBIS laser | Laser 0 | LX 637-140C | Coherent | 1 | |
| | | Laser 1 | LX 488-50C | | | |
| | | Laser 2 | LX 561-50 | | | |
| | | Laser 3 | LX 405 | | | |
| 2 | 613 nm Dichroic beam splitter | DM1 | LM01-613-25 | Semrock | 1 | |
| 3 | 503 nm Dichroic beam splitter | DM2 | LM01-503-25 | Semrock | 1 | |
| 4 | 445 nm Dichroic beam splitter | DM1 | DMLP445 | Thorlabs | 1 | |
| 5 | f = 50 mm, Ø1" achromatic doublet | L1, L2, L5, L8 | AC254-50-A-ML | Thorlabs | 4 | |
| 6 | f = 150 mm, Ø1" achromatic doublet | L3, L4 | AC254-150-A-ML | Thorlabs | 2 | |
| 7 | f = 100 mm, Ø2" achromatic doublet | L11 | AC508-100-A-ML | Thorlabs | 1 | |
| 8 | f = 100 mm, Ø1" achromatic doublet | L6 | AC254-100-A-ML | Thorlabs | 1 | |
| 9 | f = 200 mm, Ø2" achromatic doublet | L7 | ACT508-200-A-ML | Thorlabs | 1 | |
| 10 | f = 175 mm, Ø1" achromatic doublet | L10 | 49-363 | Edmund Optics | 1 | |
| 11 | f = 200 mm, Ø2" achromatic doublet | L9 | ACT508-200-A-ML | Thorlabs | 1 | |
| 12 | 50 μm pinhole | Pinhole | P50D | Thorlabs | 1 | |
| 13 | Polarizing beam splitter | PBS1 | 10FC16PB.7 | Newport Corporation | 2 | |
| 14 | f = 25 mm, Ø0.5" cylindrical achromat | CL1 | 68-160 | Edmund Optics | 1 | |
| 15 | f = 200 mm, Ø1" cylindrical achromat | CL2 | ACY254-200-A | Thorlabs | 1 | |
| 16 | f = 50 mm, Ø1" cylindrical achromat | CL3 | ACY254-50-A | Thorlabs | 1 | |
| 17 | Resonant mirror galvanometer | Resonant mirror | CRS 4 kHz | Cambridge Technology | 1 | |
| 18 | Adjustable mechanical slit | Slit | VA100 | Thorlabs | 1 | |
| 19 | Galvanometric mirror | Galvo | GVS011 | Thorlabs | 1 | |
| 20 | Quarter wave plate | QWP | AQWP3 | Bolder Vision Optik | 1 | |
| 21 | High speed focuser | FM | BLINK | Thorlabs | 1 | |
| 22 | Microscope objective (40x/NA 1.0 oil) | OBJ1 | PlanApo 40x/1.0 | Nikon | 1 | |
| 23 | Microscope objective (60x/NA 0.70 air) | OBJ2 | Dry objective | Nikon | 1 | |
| 24 | 12V DC power supply | 12V DC | A12MT400 | Acopian | 1 | |
| 25 | Quad band filter | Quad band filter | ZT 405/488/561 /640rpcv2-UF1 | Chroma | 1 | |
| 26 | Emission filter | Flip EF | FF01-525/30-25 | Semrock | 1 | |
| 27 | Emission filter | EF1 | BLP01-405R-25 | Semrock | 1 | |

| Sl. No. | Description | Item abbreviation | Part number | Part number company | Qty | Remarks |
|---|---|---|---|---|---|---|
| 28 | Emission filter | EF2 | FF01-525/45-25 | Semrock | 1 | |
| 29 | Emission filter | EF3 | FF01-593/40-25 | Semrock | 1 | |
| 30 | Emission filter | EF4 | BLP01-647R-25 | Semrock | 1 | |
| 31 | 1" x 1" protected silver mirror | M | PFSQ10-03-P01 | Thorlabs | few | |
| 32 | 2" x 2" protected silver mirror | M | PFSQ20-03-P01 | Thorlabs | few | |
| 33 | Ø1" protected silver mirror | M | PF10-03-P01 | Thorlabs | few | |
| 34 | Ø7.0 mm protected silver mirror | FM | PF03-03-P01 | Thorlabs | 1 | |
| 35 | single axis translation stage | Lin Tran stage | LT1 | Thorlabs | few | |
| 36 | 6 position motorized filter wheel | Wheel | FW102C | Thorlabs | 1 | |
| 37 | Digital sCMOS camera | Camera | Orca Flash 4.0, V2<br>model: C13440-20CU | Hamamatsu Corporation | 1 | |
| 38 | Self-Contained XYZ translation stage | 3D Tran stage | LX30 | Thorlabs | 2 | |
| 39 | 3D motorized stage | Mot stage | Model: MP-285A, PCIe 80 7852R | National Instruments | 1 | |

Qty. quantity

**Supplementary Table 3 |** *C. elegans* straining and target genotype.

| Strain | Genotype | Source |
|---|---|---|
| PHX3691 | *sqt-3(syb3691[SQT-3::mNeonGreen]) V* | S.K. Birnbaum et al.[10] |
| PON0002 | wwSi15[Pidh-1::idh-1(R156C); unc-119(+) II]; sqt-3(syb3691[SQT-3::mNeonGreen]) V | This work |
| VT1367 | maIs105 [COL-19::GFP] V | Z. Ren et al.[11] |
